# Differential Regulation of Hepatic Macrophage Fate by Chi3l1 in MASLD

**DOI:** 10.1101/2025.05.06.652369

**Authors:** Jia He, Bo Chen, Weiju Lu, Xiong Wang, Ruoxue Yang, Chengxiang Deng, Xiane Zhu, Keqin Wang, Lang Wang, Cheng Xie, Rui Li, Xiaokang Lu, Ruizhi Yang, Cheng Peng, Canpeng Li, Zhao Shan

## Abstract

Metabolic dysfunction-associated steatotic liver disease (MASLD) progression involves the replacement of protective embryo-derived Kupffer cells (KCs) by inflammatory monocyte-derived macrophages (MoMFs), yet the regulatory mechanisms remain unclear. Here, we identify chitinase 3-like 1 (Chi3l1/YKL-40) as a critical metabolic regulator of hepatic macrophage fate. We observed high expression of Chi3l1 in both KCs and MoMFs during MASLD development. Genetic deletion of Chi3l1 specifically in KCs significantly exacerbated MASLD severity and metabolic dysfunction, whereas MoMF-specific Chi3l1 deletion showed minimal metabolic effects. Mechanistic studies revealed that this cell type-specific regulation arises from differential metabolic requirements: KCs display elevated glucose metabolism compared to MoMFs. Chi3l1 directly interacts with glucose to inhibit its cellular uptake, thereby selectively protecting glucose-dependent KCs from metabolic stress-induced cell death while having negligible effects on less glucose-dependent MoMFs. These findings uncover a novel Chi3l1-mediated metabolic checkpoint that preferentially maintains KCs populations through glucose metabolism modulation, providing important new insights into the pathogenesis of MASLD and potential therapeutic strategies targeting macrophage-specific metabolic pathways.

## Introduction

Metabolic dysfunction-associated steatotic liver disease (MASLD) has become the most prevalent chronic liver disorder in western populations, affecting approximately 30% of adults and driven by its strong association with obesity and metabolic syndrome^1^. The disease spectrum ranges from matabolic dysfunction-associated fatty liver (MAFL) to metabolic dysfunction-associated steatohepatitis (MASH), with the latter characterized by steatosis, inflammation, hepatocyte ballooning, and progressive fibrosis^2^. Central to MASLD pathogenesis are hepatic macrophages, particularly the embryo-derived Kupffer cells (KCs) that reside in liver sinusoids^3,4^. These self-renewing resident macrophages^5^, play crucial roles in lipid homeostasis, as evidenced by studies showing that depletion of CD207^+^ KCs leads to impaired triglyceride storage^6^. As MASH progresses, dying KCs are progressively replaced by monocyte-derived macrophages (MoMFs) that exhibit heightened inflammatory properties and contribute to liver damage^6,7^. For example, one study demonstrated that in diet-induced MASH, KCs enhancer landscapes and gene expression profiles are profoundly reprogrammed (including up-regulation of Trem2 and Cd9) and KCs identity is lost, while MoMFs adopt convergent epigenomes, transcriptomes and functions during macrophage recruitment and adaptation in MASH^8^. Another work showed that in MASLD the number of resident KCs declines and MoMFs accumulate; these recruited macrophages include subsets that either mirror homeostatic KCs or resemble lipid-associated macrophages (LAMs) from obese adipose tissue, with the LAM-type expressing osteopontin and localizing to fibrotic zones^9^. Together, these findings highlight that this transition from protective embryo-derived KCs (EmKCs) to monocyte-derived KCs (MoKCs) represents a critical juncture in disease progression, yet the mechanisms regulating this shift remain poorly understood.

A key determinant of macrophage function is cellular metabolism. Macrophages dynamically switch between glycolytic and oxidative phosphorylation pathways to adapt to environmental changes^10^. During MASLD, hepatic macrophages increase their glycolytic activity, which may exacerbate inflammation and tissue damage^11–13^. While glucose metabolism is known to influence macrophage polarization, its specific role in determining hepatic macrophage fate - particularly the balance between KCs and MoMFs - remains unknown. Chitinase 3-like 1 (Chi3l1/YKL-40) has emerged as an important regulator of macrophage biology, promoting cell survival through ERK1/2 and PI3K/Akt pathways while modulating anti-inflammatory cytokines like IL-10^14–18^. However, its potential role in macrophage metabolic reprogramming, particularly in the context of hepatic glucose metabolism, has not been explored.

In this study, we identify a novel mechanism by which Chi3l1 governs hepatic macrophage fate through metabolic regulation. We demonstrate that Chi3l1 directly interacts with glucose to suppress its uptake in macrophages. Strikingly, this interaction selectively protects glucose-high KCs from cell death in MASLD conditions, while having minimal effect on glucose-low MoMFs. These findings reveal a previously unrecognized Chi3l1-mediated metabolic checkpoint that maintains KC populations, providing new insights into the pathogenesis of MASLD and potential therapeutic strategies.

## Materials and methods

### Animal experiments and procedures

*Animals Chil1^-/-^*(strain no. T014402), *Chil1^flox//flox^* (strain no. T013652), *Lyz2-cre* (strain no. T003822), *Clec4f-cre* (strain no. T036801) with a *C57BL/6J* background were purchased from GemPharmatech. *Rosa tdtomato* mice (strain no. C001181) were purchased from Cyagen. Accordingly, *C57BL/6J* mice (strain no. N000013) were used as wild-type (WT) mice. To generate *Clec4f^△Chil^*^1^ mice, *Chil1^flox//flox^* mice were crossed with *Clec4f-cre* mice. To generate *Clec4f^Rosa^ ^tdtomato^* mice, *Rosa tdtomato* mice were crossed with *Clec4f-cre* mice. To generate *Lyz2^△Chil^*^1^ mice, *Chil1^flox//flox^*mice were crossed with *Lyz2-cre* mice. Male mice have been the choice in the vast majority of the studies of MASLD reported in the literature^19,20^. Therefore, we used male mice in the majority of the experiments presented. All mouse colonies were maintained at the Animal Core Facility of Yunnan University. The animal studies were approved by the Yunnan University Institutional Animal Care and Use Committee (IACUC, Approval No. YNU20220314).

*Construction of MASLD/MASH mouse model* Mice were provided a high-fat and high-cholesterol diet (Research Diet, d12108c, 40 kcal% fat and 1.25% cholesterol) for 16 weeks or a methionine and choline deficient diet (Research Diet, A02082002BR) for 6 weeks. Throughout the feeding period, the body weight and food consumption of the mice were observed and recorded weekly. Once the dietary intervention was completed, the mice were euthanized. Liver and murine serum samples were collected for further analysis. Alanine aminotransferase (ALT) and aspartate aminotransferase (AST) levels in the serum, as well as cholesterol (TC) and triglyceride (TG) levels in both serum and liver tissues, were quantified using commercially available kits (Nanjing Jiancheng Bioengineering Institute).

### Statistical analysis

Data are presented as mean ± standard error of the mean (SEM) in all graph figures. Statistical analyses were conducted using the SPSS statistics software (Version 22). To compare the two groups, an unpaired two-tailed Student’s t-test was used. One-way analysis of variance (ANOVA) was performed for comparisons involving three or more groups. For patients with MASLD liver, the samples were tested using the Mann-Whitney test. Statistical significance was set at p < 0.05 and p value is indicated. All cell culture results represent at least three independent experiments.

### Additional Methods

Additional detailed methods can be found in the Supporting Information.

## Results

### Hepatic macrophages express Chi3l1 and upregulate its expression post HFHC diet

In our previous study, we found that Chi3l1 expressed by hepatic macrophages influences macrophage function during acute liver injury^21^. Therefore, we sought to determine whether Chi3l1 also plays a role in MASLD and whether its expression in hepatic macrophages is altered in this context. To this end, we first established a mouse model of MASLD by feeding C57BL/6J wild-type mice a normal chow diet (NCD) or an high-fat high-cholesterol (HFHC) diet for 16 weeks. Histological analysis of liver sections using Hematoxylin and Eosin (H&E) and Sirius Red staining revealed marked lipid accumulation without apparent fibrosis (Figure S1A). Consistently, western blot analysis showed no upregulation of α-SMA (Figure S1B), suggesting that our HFHC model represents an early stage of MASLD.

Next, we performed immunofluorescence staining for Chi3l1 in liver sections using antibodies against TIM4 (a KCs marker), F4/80 (a pan-macrophage marker), and Chi3l1. This confirmed that Chi3l1 is expressed in both KCs (TIM4^+^F4/80^+^ cells) and MoMFs (TIM4^-^F4/80^+^ cells), with elevated expression under HFHC conditions (Figure 1A). To validate the specificity of Chi3l1 staining, we generated *Chil1^-/-^* mice and confirmed knockout efficiency by qRT-PCR (Figure S2A, B). Immunofluorescence staining for Chi3l1 in liver sections from WT and *Chil1^-/-^* mice showed that the anti-Chi3l1 antibody specifically detected Chi3l1 in WT but not *Chil1^-/-^* mice, confirming the specificity of the staining (Figure 1B). We next assessed whether Chi3l1 is upregulated by HFHC feeding by measuring its protein levels in isolated KCs and whole liver tissue via western blotting. A marked increase in Chi3l1 expression was observed in both KCs and liver tissue following HFHC diet feeding (Figure 1C, D). Consistently, patients with MAFL or MASH exhibited elevated hepatic Chi3l1 mRNA levels, which correlated with MASLD severity and fibrosis stage (Figure 1E, F). These findings suggest that Chi3l1 is expressed in hepatic macrophages and may contribute to MASLD progression.

**Figure 1.**
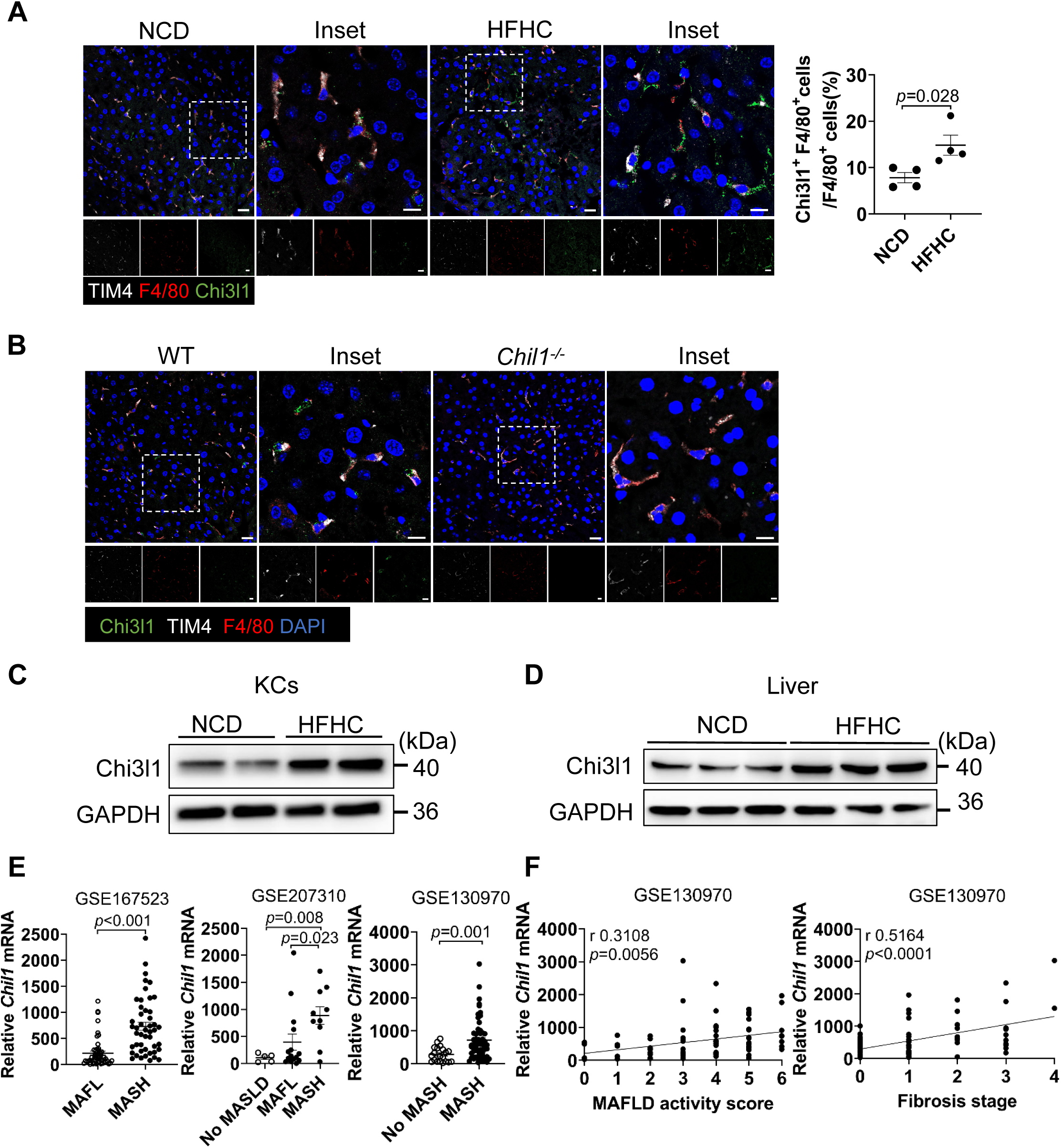
Hepatic macrophages express Chi3l1 and upregulate its expression post HFHC diet. **(A)** Immunofluorescent staining of TIM4 (white), F4/80 (red), Chi3l1 (green), and nuclear DAPI (blue) in liver sections of mice fed with either NCD or HFHC for 16 weeks, illustrating Chi3l1 expression in hepatic macrophages. Scale bar=20 µm and 10 µm (Inset). Chi3l1^+^ F4/80^+^ cells/F4/80^+^ cells were statistically analyzed. n=4 mice/group. **(B)** Representative immunofluorescence images of liver sections from WT and *Chil1^-/-^* mice stained for Chi3l1(green), F4/80 (macrophages), and TIM4 (Kupffer cells). DAPI (blue) marks nuclei. Scale bar=20 µm and 10 µm (Insets). **(C, D)** Western blot analysis of Chi3l1 in either isolated Kupffer cells (KCs, F) or whole liver tissue (Liver, G) from mice fed either NCD or HFHC diet. n=2-3 mice/group. **(E)** mRNA expression levels of Chil1 in liver tissues of patients with metabolic dysfunction-associated fatty liver (MAFL) or with metabolic dysfunction-associated steatohepatitis (MASH) (GEO Datasets: GSE167523, GSE207310, GSE130970). No-MAFLD or Healthy individuals serve as controls. **(F)** The correlation between mRNA expression levels of Chil1 and MASLD activity score or fibrosis stage was analyzed (GEO Datasets: GSE130970). Representative images were shown in A, B. Mann-Whitney test was performed in E. Pearson’s correlation was performed in F. P value and r value are as indicated.

### Deficiency of Chi3l1 in Kupffer cells promotes insulin resistance and hepatic lipid accumulation

Given that Chi3l1 is highly expressed in hepatic macrophages, we investigated its functional role by generating mice with conditional knockout (cKO) of *Chil1* in either KCs or MoMFs. First, we generated *Clec4f^ΔChil^*^1^ mice by crossing *Chil1^fl/fl^* mice with *Clec4f-cre* mice^22^, achieving KC-specific deletion of Chil1 (Figure S3A-C). These mice, along with *Chil1^fl/fl^* controls, were fed either a NCD or a HFHC diet. Under NCD feeding, *Clec4f^ΔChil^*^1^ and *Chil1^fl/fl^* mice displayed comparable phenotypes in terms of body weight gain, hepatic lipid deposition, metabolic parameters, glucose tolerance and insulin resistance (Figure 2A–F). In contrast, when fed an HFHC diet, *Clec4f^ΔChil^*^1^ mice exhibited markedly accelerated weight gain compared to controls (Figure 2A, B). These mice also showed increased hepatic lipid accumulation, as evidenced by H&E and Oil Red O staining at 16 weeks (Figure 2C), along with greater metabolic disturbances, including a higher liver index (liver-to-body weight ratio), elevated serum ALT levels, and increased cholesterol and triglyceride levels in both liver and serum (Figure 2D). Furthermore, *Clec4f^ΔChil^*^1^ mice exhibited impaired glucose metabolism, as indicated by worsened glucose tolerance and insulin resistance in IGTT and ITT assays (Figure 2E, F). To exclude potential off-target effects caused by *Clec4f-Cre* insertion, we compared *Clec4f-Cre* and *Clec4f^ΔChil^*^1^ mice. The phenotypic differences between *Clec4f-Cre* and *Clec4f^ΔChil^*^1^ mice mirrored those observed between *Chil1^fl/fl^* and *Clec4f^ΔChil^*^1^ mice, with the latter showing faster weight gain, more severe hepatic steatosis, greater metabolic dysregulation, and worsened glucose intolerance and insulin resistance (Figure S4A–G).

**Figure 2.**
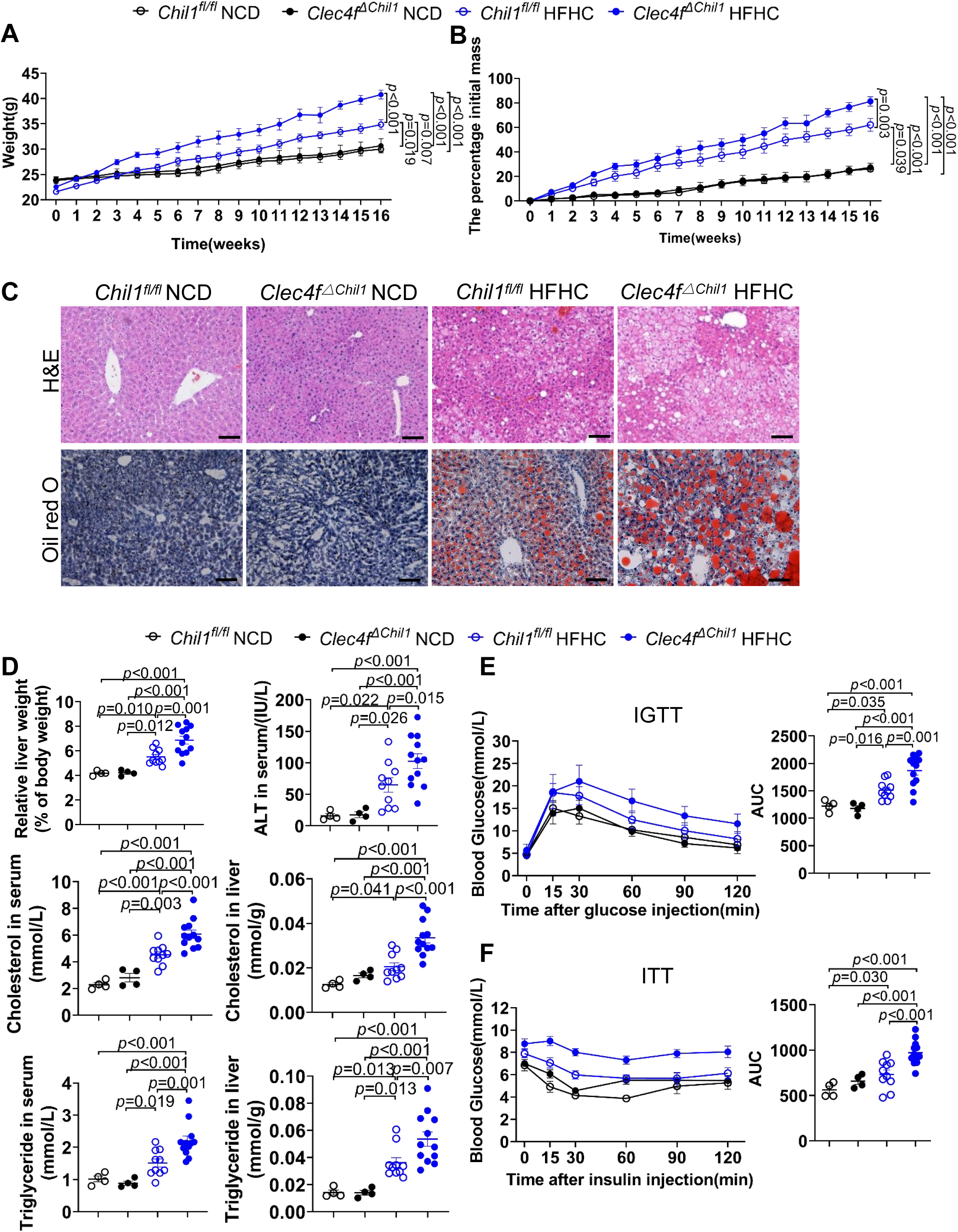
Deficiency of Chi3l1 in Kupffer cells promotes insulin resistance and hepatic lipid accumulation. *Chil1^fl/fl^* and *Clec4f^ΔChil^*^1^ mice were fed either a normal chow diet (NCD) or a high-fat, high-cholesterol (HFHC) diet for 16 weeks. **(A, B)** Body weight was recorded during HFHC diet feeding (A) and expressed as a percentage of initial body mass (B). **(C)** H&E (Upper panel) and oil red o staining (Lower panel) was performed to examine liver histology and hepatic lipid accumulation in both genotypes after 16 weeks of NCD or HFHC diet. Scale bar = 20 µm. **(D)** Liver index (liver weight/body weight × 100%), ALT levels, and serum and liver Cholesterol or Triglyceride levels were measured in both genotypes after 16 weeks on NCD or HFHC diets. n=4-12 mice/group. **(E, F)** Intraperitoneal glucose tolerance test (IGTT) and insulin tolerance test (ITT) were performed after 16 weeks of NCD or HFHC feeding in both genotypes (n = 4–12 mice per group). Representative images were shown in (C). One-way ANOVA was performed in (A, B, D-F). P-value is as indicated.

To investigate the role of KCs-derived Chi3l1 in MASH, we first examined its expression in a methionine-choline deficient (MCD) diet model. Wildtype mice fed an MCD diet for 6 weeks showed significantly increased Chi3l1 mRNA and protein levels in whole liver tissues compared to NCD controls, confirming diet-induced upregulation (Figure 3A, B). To assess the functional contribution of KCs-derived Chi3l1, we subjected *Clec4f^ΔChil^*^1^ mice along with *Chil1^fl/fl^*controls to 6 weeks of MCD diet feeding. Body weight was comparable between genotypes throughout the feeding period (Figure 3C). Histological analysis revealed that loss of Chi3l1 in Kupffer cells led to a significant exacerbation of MCD diet-induced hepatic steatosis, inflammation, and fibrosis, as reflected by increased MASLD activity scores, Oil Red O staining, Sirius Red deposition, and α-SMA expression (Figure 3D). Consistent with this histological finding, *Clec4f^ΔChil^*^1^ mice exhibited an increased liver index but similar serum ALT levels, reflecting liver weight gain without evidence of enhanced liver injury (Figure 3E). Additionally, these mice showed significant increases in serum and hepatic triglyceride levels, as well as elevated serum cholesterol, whereas hepatic cholesterol is not significantly upregulated compared to controls (Figure 3E). These data demonstrate that loss of Chi3l1 in Kupffer cells promotes hepatic steatosis, suggesting a protective role for KC-derived Chi3l1 in MASH pathogenesis.

**Figure 3.**
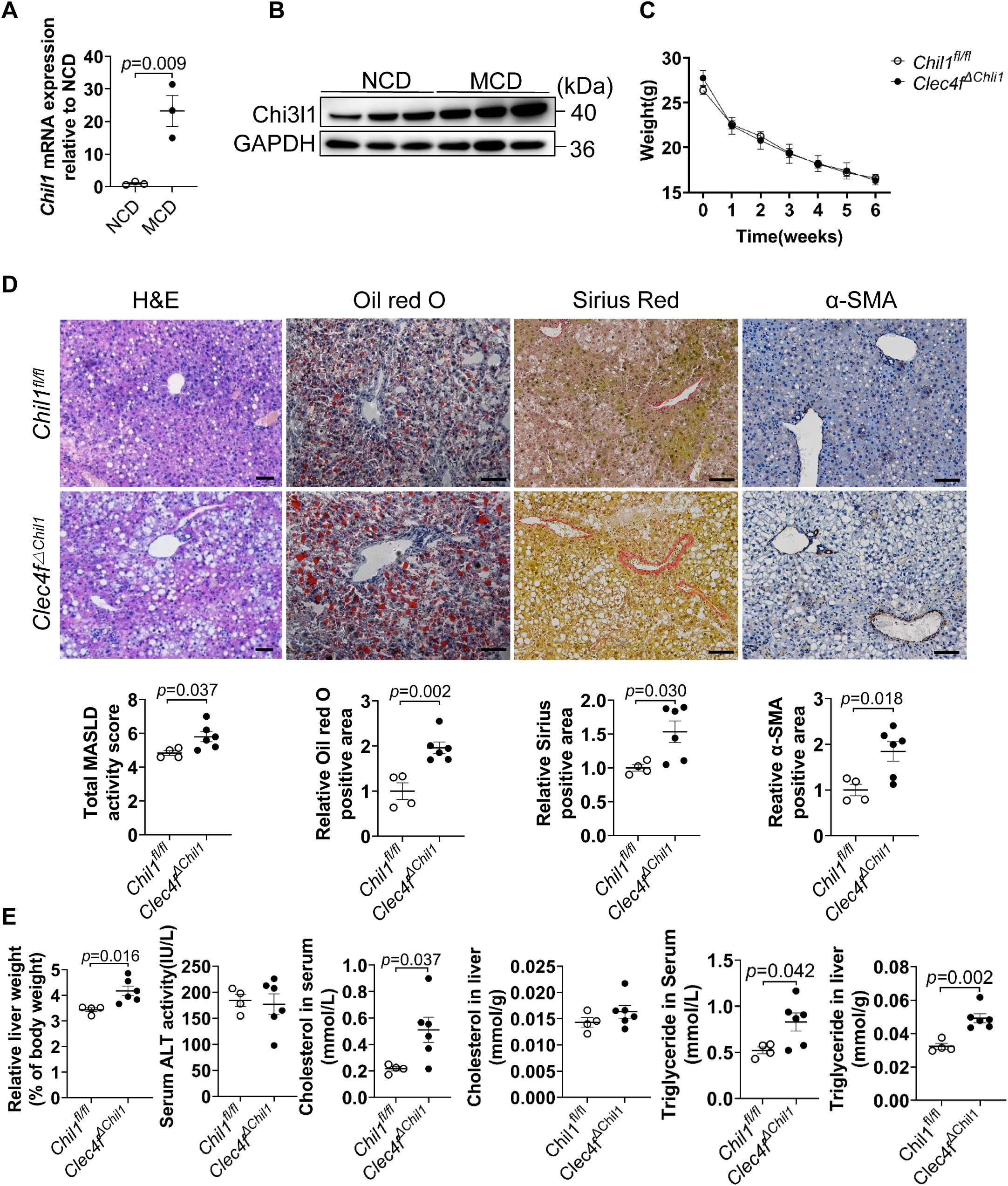
Deficiency of Chi3l1 in Kupffer cells promotes liver steatosis and fibrosis in MASH. Male wildtype C57B/6J mice were fed with NCD or MCD diet for 6 weeks (A-B). *Chil1^fl/fl^* and *Clec4f^ΔChil^*^1^ mice were fed with a MCD diet for 6 weeks (C-E). **(A, B)** qRT-PCR (A) and wetern blot (B) analysis of Chi3l1 expression in whole liver tissues under NCD and MCD diets. n=3 mice/group. **(C)** Body weight of mice with conditional deletion of *Chil1* in KCs (*Clec4f^ΔChil^*^1^) and their control mice (*Chil1^fl/fl^*) was recorded during MCD diet. **(D)** Histological analyses were performed in liver tissue of *Clec4f^ΔChil^*^1^ and *Chil1^fl/fl^*fed the MCD diet for 6 weeks. Scale bar=20μm. **(E)** Liver index (liver weight/body weight × 100%), ALT levels and serum and liver Cholesterol or Triglyceride levels were measured in both genotypes fed the MCD diet for 6 weeks. n=4-6 mice/group. Representative images are shown in D. Two-tailed, unpaired student t-test was performed in A, C, D, E. P value is as indicated.

To assess the role of Chi3l1 in MoMFs, we generated *Lyz2^ΔChil^*^1^ mice (*Chil1^fl/fl^* × *Lyz2-Cre*^22^) and validated Chil1 deletion efficiency in MoMFs and BMDM (Figure S5A–C). Chil1 expression was completely abolished in MoMFs from *Lyz2^ΔChil^*^1^ mice. Considering the partial activity of *Lyz2-Cre* in KCs, we further assessed Chi3l1 expression in KCs isolated from *Lyz2^Chil^*^1^ mice. Only a modest (∼40%) reduction in Chi3l1 mRNA and protein levels was observed in KCs, indicating that *Lyz2-Cre*–mediated deletion minimally affects Chi3l1 expression in KCs (Figure S5B–C). *Lyz2^ΔChil^*^1^ and *Chil1^fl/fl^* control mice were then fed either a NCD or a HFHC diet. Under both dietary conditions, the two genotypes exhibited comparable phenotypes with respect to body weight gain, hepatic lipid accumulation, metabolic parameters, glucose tolerance, and insulin sensitivity (Figure S6A–F). These results indicate that Chi3l1 loss in MoMFs does not substantially impact metabolic regulation.

### ScRNA-seq reveals upregulated glucose metabolism-related transcripts in KCs, correlating with cell death signatures

To dissect the distinct metabolic and functional profiles between KCs and MoMFs during MASLD progression, we performed BD Rhapsody single-cell RNA sequencing (scRNA-seq) on liver non-parenchymal cells (NPCs) from mice fed a NCD or HFHC diet for 16 weeks. After quality control and filtration, we retained 23,312 cells from NCD livers and 6,567 cells from HFHC livers for downstream analysis. Using a graph-based clustering approach, we identified 32 distinct cell populations, visualized via uniform manifold approximation and projection (UMAP) (Figure 4A). Monocyte/macrophage subsets were further defined based on lineage-specific markers: Monocytes expressed *Ly6c2, Chil3, S100a6, Ccr2, Itgam*, and *Cx3cr1* but lacked macrophage markers. KCs were marked by *Cd68, Vsig4, Clec4f, TIM4, Adgre1*, and *Clec1b*. MoMFs were negative for KCs markers but positive for macrophage markers such as *Ccr2, Cx3cr1, Cd9, Itgax, Gpnmb, Cd68,* and *Adgre1* (Figure 4B, C; UMAP in Figure S7A)^8,23^.

**Figure 4.**
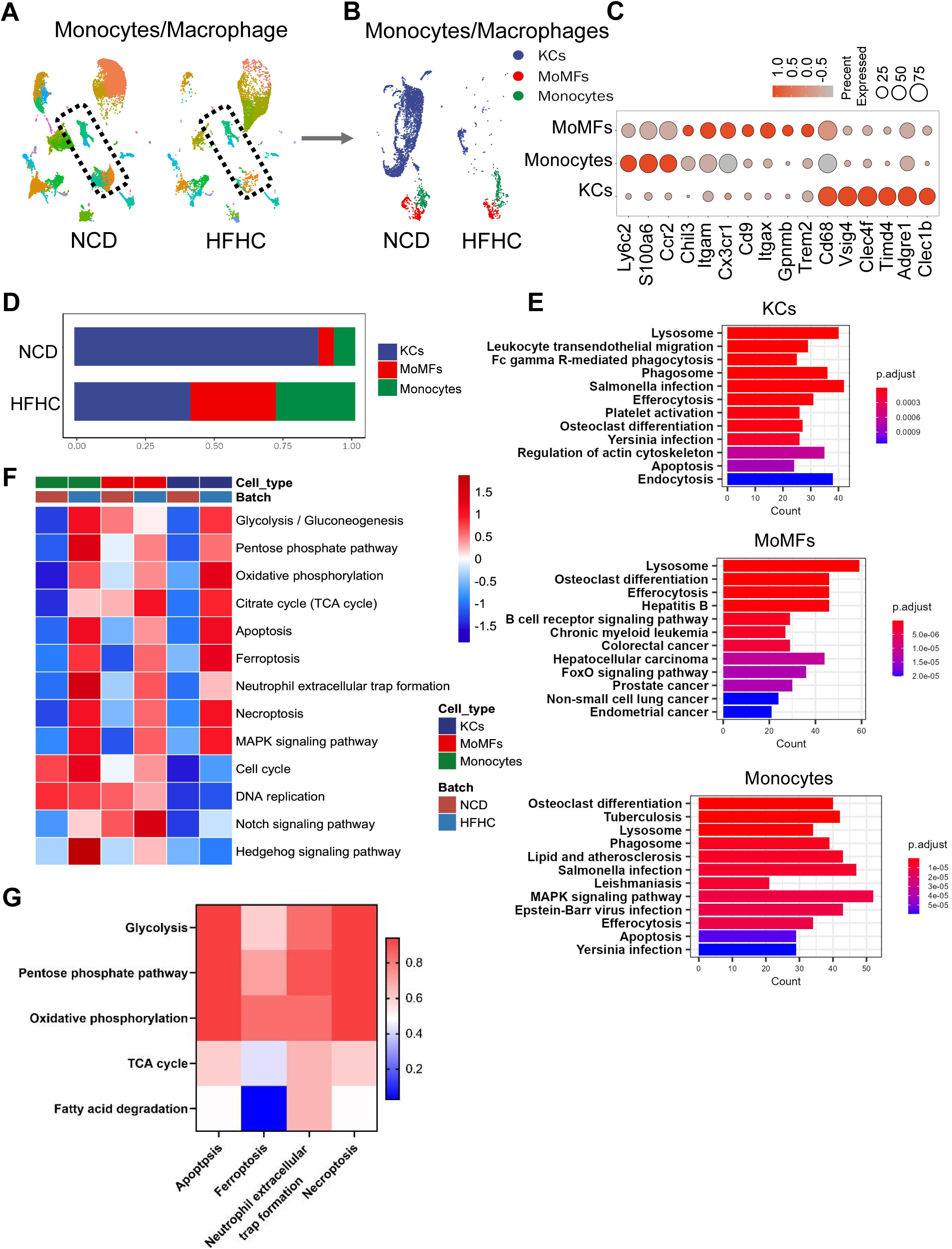
ScRNA-seq reveals upregulated glucose metabolism-related transcripts in KCs, correlating with cell death signatures. Wildtype C57BL/6J mice were fed either a normal chow diet (NCD) or HFHC for 16 weeks. NPCs were isolated and subjected to BD Rhapsody scRNA sequencing. **(A)** Uniform manifold approximation and projection (UMAP) plots illustrate the clustering of NPCs in the livers of mice fed NCD and HFHC. Cell clusters are color-coded, with monocytes/macrophages clusters outlined. **(B)** UMAP plots depict the clustering of Monocytes/Macrophages in the livers of mice fed NCD and HFHC. Cell clusters are color-coded. **(C)** Dot plot displays the scaled gene expression levels of lineage-specific marker genes in different cell clusters. **(D)** Quantification of each cell cluster is presented. **(E)** KEGG analysis reveals the top 12 enriched pathways for up-regulated genes when comparing HFHC versus NCD in KCs, monocytes, and MoMFs, respectively. **(F)** Gene set variation analysis (GSVA) shows pathway activity for cell death, glucose metabolism, and cell proliferation in KCs, monocytes, and MoMFs of WT mice fed NCD or HFHC for 16 weeks, respectively. **(G)** The correlation between cell death and glucose metabolism pathways, based on GSVA score, is depicted.

Consistent with prior studies^6,7^, we observed decreased KCs numbers but increased MoMFs and monocytes in HFHC-fed mice compared to NCD controls (Figure 4D). Kyoto Encyclopedia of Genes and Genomes (KEGG) pathway analysis showed that while both cell types exhibited activation of phagocytosis-related pathways (lysosome, phagosome, endocytosis, and efferocytosis), they displayed divergent cell fate patterns (Figure 4E). KCs showed strong cell death signatures, whereas MoMFs maintained proliferative activity without evidence of cell death (Figure 4E). Monocytes showed strong cell death and proliferative activity (Figure 4E). Given the significant role of metabolic regulation in cell fate,^24^ we compared pathways involved in glucose metabolism, cell death, and cell proliferation (Figure 4F). Notably, glucose metabolism pathways were significantly more active in KCs and monocytes compared to MoMFs (Figure 4F). Moreover, the cell proliferation pathway was highly activated in monocytes and consistently activated in MoMFs but not in KCs (Figure 4F). Gene Set Variation Analysis (GSVA)-based correlation analysis revealed a striking association between glucose metabolism and cell death pathways (Figure 4G). These findings demonstrate distinct glucose metabolic activation patterns between KCs and MoMFs, which may underlie their divergent cell fates in MASLD progression.

### Chi3l1 deficiency promote KCs death during MASLD

To investigate the role of Chi3l1 in KCs survival during MASLD, we then performed scRNA-seq on NPCs isolated from *Chil1^-/-^* mice fed an HFHC diet for 16 weeks. After quality control, 6,813 high-quality cells were retained for analysis. Using established KC markers (Figure 4C), we conducted GSVA to examine metabolic pathways. This revealed enhanced cell death pathways in KCs from HFHC-fed mice, with significantly greater apoptosis signatures in *Chil1^-/-^* KCs compared to wild-type (WT) controls (Figure 5A). The increased apoptosis was further supported by upregulation of pro-apoptotic genes in *Chil1^-/-^*KCs (Figure 5B).

**Figure 5.**
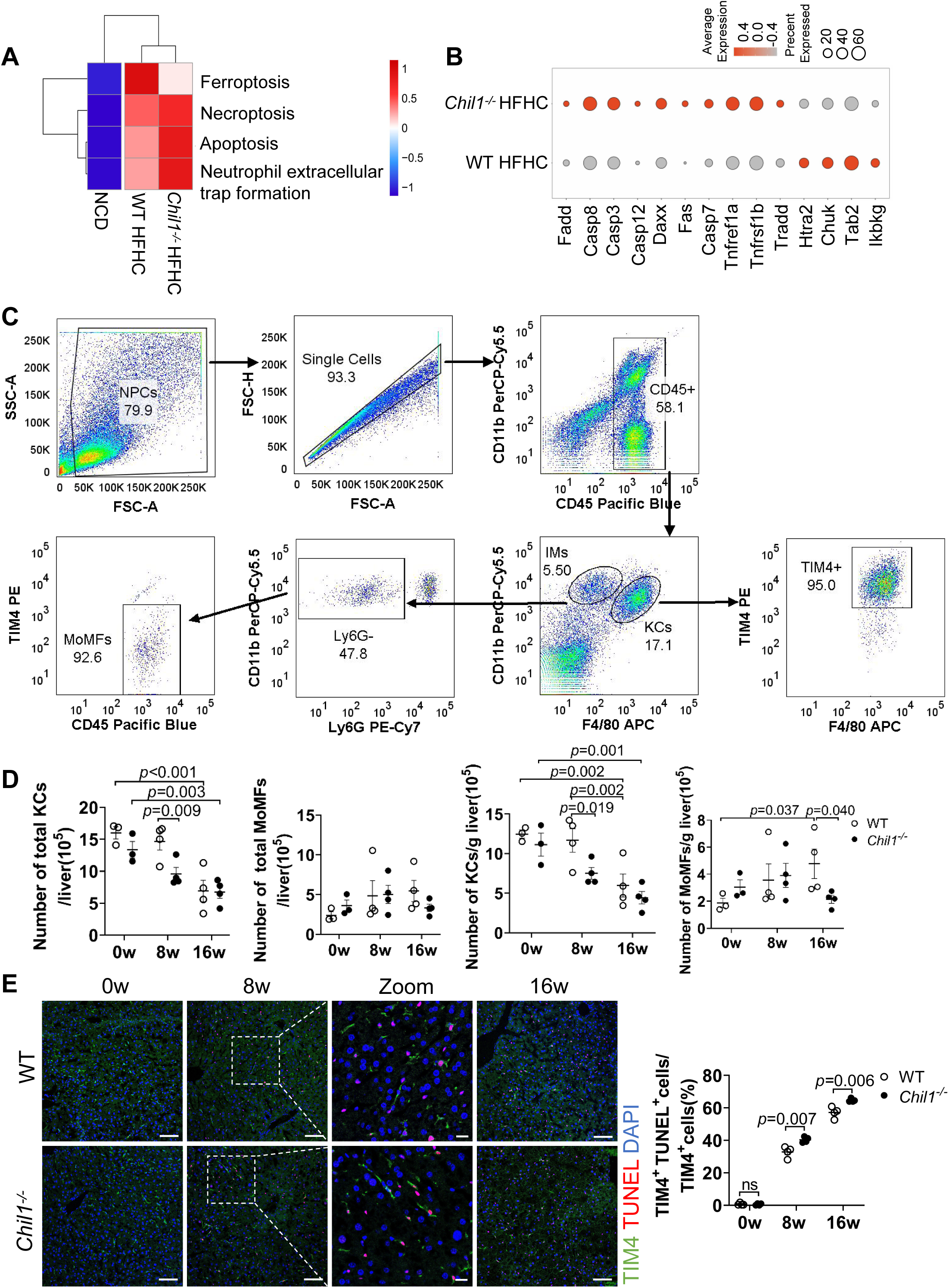
Chi3l1 deficiency promote KCs death during MASLD. **(A)** GSVA analysis showed the enrichment of cell death-related pathways in KCs from WT mice fed with either NCD or HFHC or *Chil1^-/-^* mice fed with HFHC. **(B)** Dot plot showing the scaled gene expression levels of Apoptosis-related genes and repressor genes in KCs from either WT or *Chil1^-/-^*fed with HFHC. **(C)** Strategy used to gate KCs (CD45^+^ F4/80^hi^ CD11b^low^ TIM4^hi^) and MoMFs (CD45^+^ F4/80^low^ CD11b^hi^ Ly6G^-^ TIM4^-^) in the liver by flow cytometry. **(D)** Number of KCs and MoMFs /liver or gram(g) liver were statistically analyzed. n= 3-4 mice per group. **(E)** Immunofluorescent staining to detect TIM4(green), TUNEL (red), and nuclear DAPI (blue) in liver sections. Scale bar=50µm and 20µm (Insets). TUNEL^+^ TIM4^+^ cells/TIM4^+^ cells were statistically analyzed. n=4 mice/group. Representative images are shown in C, E. One-way ANOVA was performed in D. Two-tailed, unpaired student t-test was performed in E. P value is as indicated.

We next validated these findings by flow cytometry using the gating strategy shown in Figure 5C. While WT and *Chil1^-/-^* mice showed similar KC numbers at baseline, dramatic differences emerged during HFHC feeding. WT KC numbers remained stable at 8 weeks but decreased by 50% at 16 weeks. In contrast, *Chil1^-/-^* mice exhibited accelerated KCs loss, with a 30% reduction by 8 weeks progressing to 60% by 16 weeks (Figure 5D, S8A). Notably, MoMFs populations remained comparable between groups at early timepoints but showed greater reduction in *Chil1^-/-^* mice at 16 weeks (Figure 5D, S8A).

Histological analysis further supported these findings. TIM4/TUNEL co-staining revealed no TUNEL⁺ KCs in WT livers at baseline, whereas 40% and 50% of KCs were TUNEL⁺ at 8 and 16 weeks, respectively. In *Chil1^-/-^* mice, KC apoptosis was significantly increased at both time points (Figure 5E). Consistent results were obtained with TIM4/cleaved caspase-3 co-staining (Figure S8B). We further confirmed these observations in *Clec4f^ΔChil^*^1^ mice in both HFHC^25^ and MCD diet models. In the MCD model, *Clec4f^ΔChil^*^1^ mice exhibited enhanced KCs death compared with *Chil1^fl/fl^* controls (Figure S9A). To exclude potential effects of myeloid cell–derived Chi3l1 on KCs survival, we compared KCs death and abundance between *Chil1^fl/fl^* and *Lyz2^ΔChil^*^1^ mice using histological and flow cytometric analyses. Loss of Chi3l1 in MoMFs did not lead to significant KC apoptosis or depletion (Figure S9B–D). Together, these results demonstrate that Chi3l1 deficiency promotes KC apoptosis, resulting in premature KC depletion during MASLD progression.

### Molecular interaction between Chi3l1 and glucose

Our investigation into Chi3l1-mediated KCs survival revealed an unexpected structural relationship: Chi3l1 binds to glucose, which is structurally analogous to chitin, a polysaccharide well known to bind Chi3l1 (Figure 6A). Bioinformatics analysis using the STITCH database further supported this observation, predicting a high probability of direct Chi3l1-glucose interaction (Figure 6B). To experimentally validate this interaction, we performed pull-down assays using biotin-labeled glucose incubated with plasma from HFHC-fed mice. Streptavidin bead isolation followed by anti-Chi3l1 Western blotting demonstrated specific binding between Chi3l1 and biotin-glucose, but not biotin alone (Figure 6C, D). This interaction was competitively inhibited by unlabeled glucose, confirming specificity (Figure 6D). Quantitative analysis using microscale thermophoresis with recombinant mouse Chi3l1 (rChi3l1) yielded a dissociation constant (Kd) of 4.95 mM for the Chi3l1-glucose interaction (Figure 6E). Notably, circulating Chi3l1 levels were significantly elevated in serum from HFHC-fed mice compared to baseline (Figure 6F), suggesting a potential physiological role for this interaction in metabolic regulation. These findings establish Chi3l1 as a novel glucose-binding protein that may participate in glucose homeostasis during MASLD progression.

**Figure 6.**
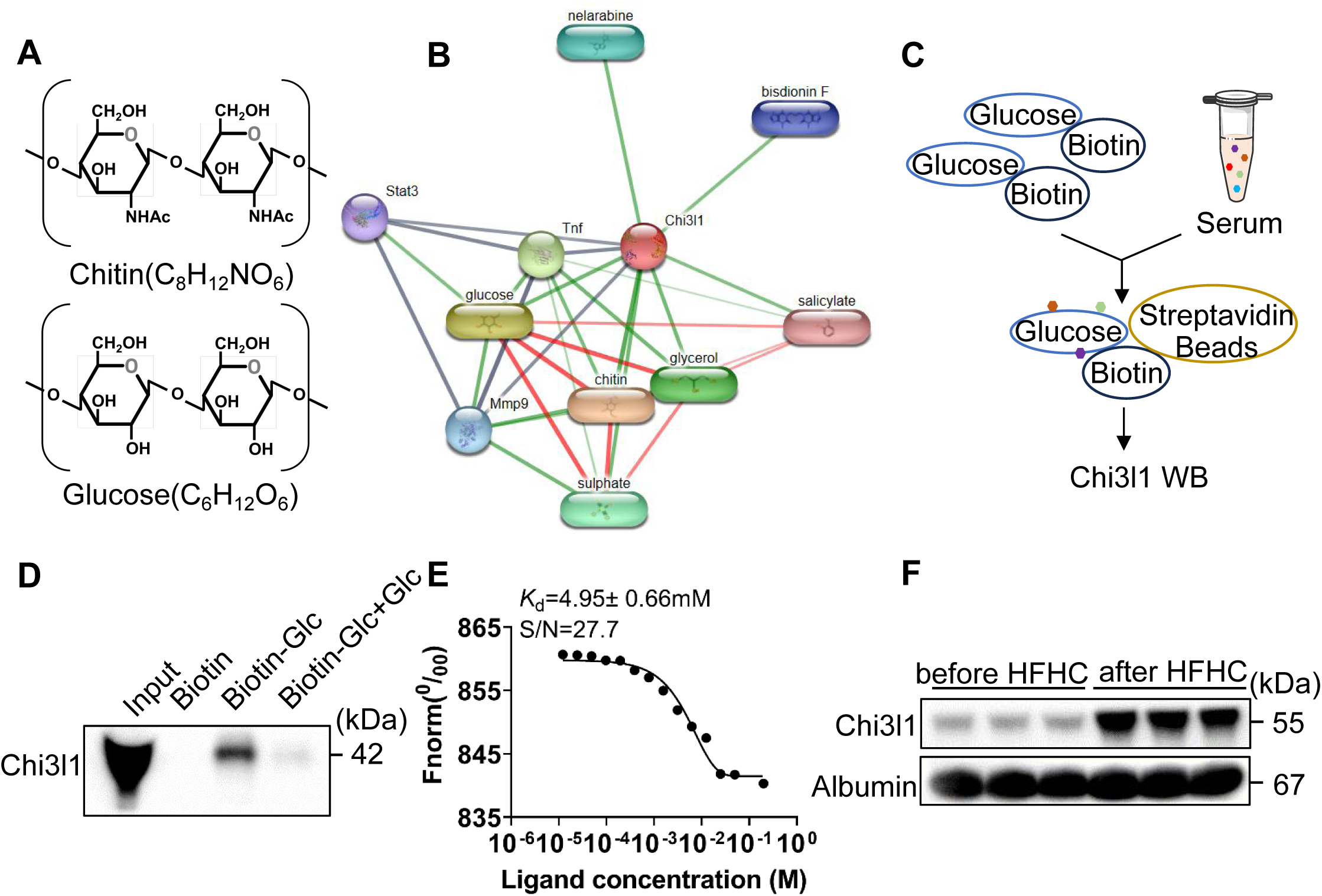
Molecular interaction between Chi3l1 and glucose. **(A)** A comparison of chemical structures between glucose and chitin. **(B)** Prediction of Chi3l1-glucose interaction using STITCH database (http://stitch.embl.de). **(C)** Strategy for pulling down glucose-binding proteins in murine serum. **(D)** Biotin-conjugated glucose was incubated with murine serum from mice fed with HFHC for 16 weeks. Proteins bound to glucose were precipitated by streptavidin beads. Biotin or biotin-conjugated glucose plus glucose were used as negative controls. Western blot was performed to examine Chi3l1 in the precipitate. **(E)** Microscale thermophoresis assay to detect the interaction between recombinant mouse Chi3l1 (rChi3l1) and glucose. Kd=4.95±0.66mM. **(F)** Western blot to detect Chi3l1 expression in murine serum before and after HFHC feeding. n=3 mice/group.

### Chi3l1 limits glucose uptake and protects hepatic macrophages from cell death

To elucidate the functional consequences of Chi3l1-glucose binding, we examined glucose metabolism in hepatic macrophages. Using the fluorescent glucose analog 2-NBDG^26^, we performed uptake assays in KCs following 12-hour glucose starvation. While glycogen droplet size remained unchanged in untreated KCs regardless of rChi3l1 supplementation (Figure 7A), 2-NBDG exposure significantly increased glycogen accumulation. This effect was markedly suppressed by rChi3l1 co-treatment (Figure 7A), a phenotype replicated in BMDM (Figure 7A). These results demonstrate that Chi3l1 restricts glucose uptake and subsequent glycogen storage.

**Figure 7.**
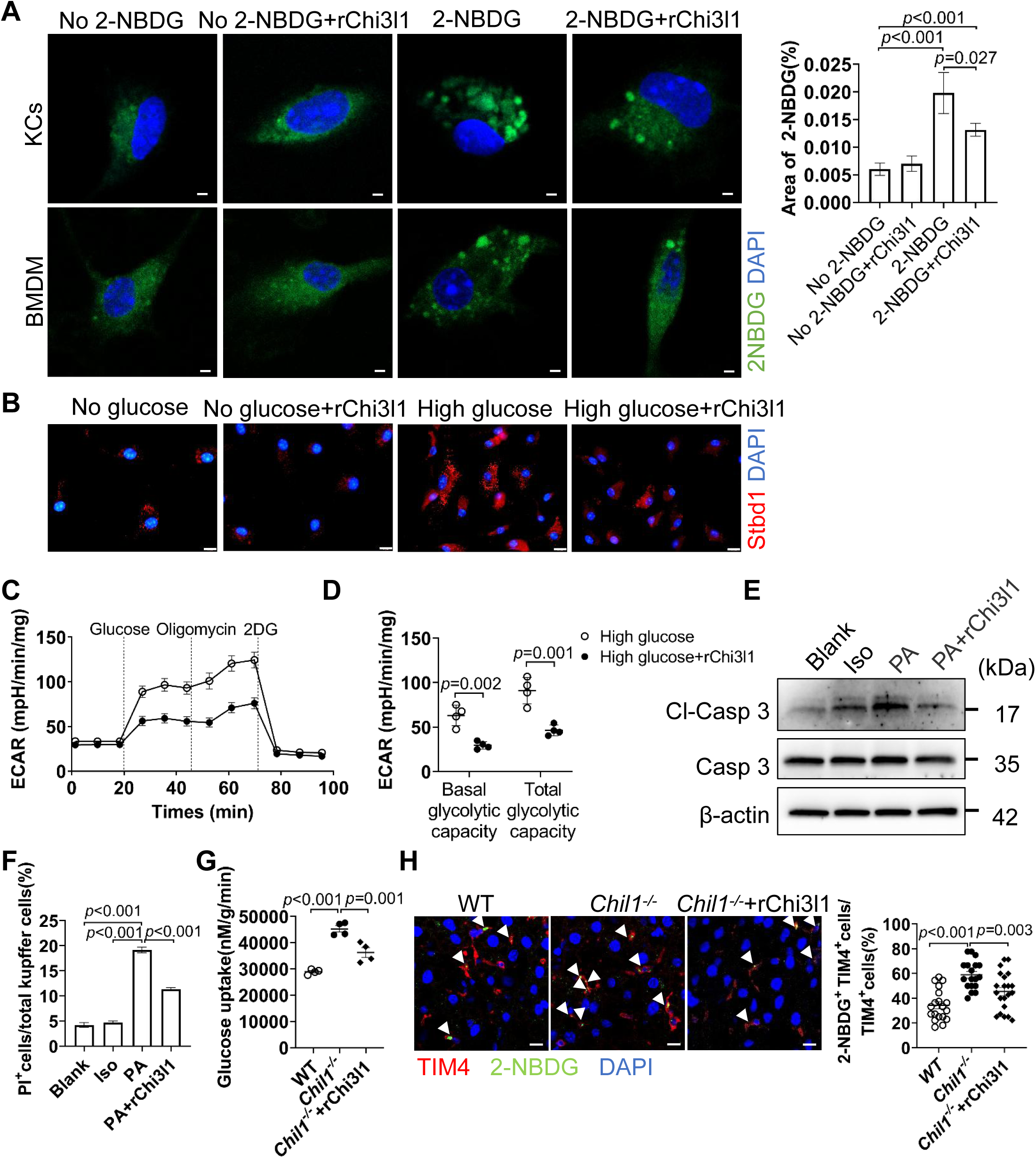
Chi3l1 limits glucose uptake and protects hepatic macrophages from cell death. **(A)** Following 12 h of glucose starvation, isolated KCs or BMDM were divided into two groups: one treated with no 2-NBDG and the other with 2-NBDG. Within each group, KCs or BMDM were further treated without or with recombinant murine Chi3l1 (rChi3l1) for 6 h. Glycogen aggregate formation labeled by 2-NBDG (Green) in KCs or BMDM was examined after counterstaining with nuclear DAPI (Blue). Scale bar=2μm. Area of 2-NBDG in KCs were quantified. **(B)** Following 12 h of glucose starvation, BMDM were treated with either no glucose or high glucose (25mM). Concurrently, BMDM were treated without or with rChi3l1 for 24 h under each condition. glycogen aggregate formation in BMDM was detected using immunofluorescence staining for Stbd1 (red) and nuclear DAPI (blue). Scale bar = 10 μm. **(C and D)** BMDM cells were treated without or with rChi3l1 for 24 h and subjected to Seahorse metabolic analysis to measure the extracellular acidification rate (ECAR). **(E and F)** KCs were treated without (blank) or with either Isopropyl alcohol (Iso) or 800uM palmitic acid (PA) or 100ng rChi3l1 with 800 uM PA for 24 h. Western blot was performed to detect cleaved caspase 3 (Cl-Casp3) in E. Calcein/PI staining was quantified to detect cell viability in F. Scale bar=50μm. **(G)** Measurement of 2-NBDG (a fluorescent glucose analog) uptake by KCs *in vivo*. WT and *Chil1^-/-^*mice, either untreated or supplemented with rChi3l1, were injected intraperitoneally with 12 mg/kg 2-NBDG. After 45mins, KCs were isolated and glucose uptake assessed by spectrophotometry. **(H)** Representative immunofluorescence images of liver sections stained for TIM4 (red) and 2-NBDG uptake (green) to visualize glucose uptake by KCs in situ. Scale bar = 10 µm (Insets). Quantification is shown as the percentage of TIM4⁺ cells that are also 2-NBDG⁺. Representative images were shown in A, B, H. One-way ANOVA was performed in A, F, G, H. Two-tailed, unpaired student t-test was performed in D. P value is as indicated.

Further validation using Stbd1 (a glycogen-binding protein^26^) immunofluorescence revealed minimal glycogen foci in glucose-deprived BMDM, with no rChi3l1-dependent differences. High-glucose conditions, however, triggered robust glycogen aggregation, which was significantly attenuated by rChi3l1 (Figure 7B). Concordantly, extracellular acidification rate (ECAR) measurements showed reduced basal and total glycolytic capacity in rChi3l1-treated BMDMs (Figure 7C, D), confirming Chi3l1’s role in limiting glucose metabolism.

To test whether Chi3l1-glucose binding influence cell survival, we employed a palmitic acid (PA)-induced lipotoxicity cell-based model to better mimic the *in vivo* environment. rChi3l1 supplementation reduced PA-induced cleavage of caspase-3 (Figure 7E) and decreased KCs death (calcein/PI staining, Figure 7F). To validate this mechanism *in vivo*, we intraperitoneally injected 2-NBDG into WT and *Chil1^-/-^* mice, with or without supplementation of rChi3l1, to assess glucose uptake by KCs. *Chil1^-/-^* KCs displayed markedly increased 2-NBDG uptake compared with WT controls, whereas rChi3l1 supplementation significantly reduced glucose uptake. These results demonstrate that serum Chi3l1 limits glucose uptake by KCs *in vivo* (Figure 7G, H). Collectively, these findings demonstrate that Chi3l1 protects KCs from metabolic stress–induced death by regulating glucose uptake.

## Discussion

Our findings establish Chi3l1 as a critical metabolic regulator that controls hepatic macrophage fate through a novel glucose-dependent mechanism in MASLD. Using cell-specific knockout models, we uncovered a fundamental dichotomy in Chi3l1 function: selective ablation in KCs dramatically accelerated MASLD progression and metabolic dysfunction, whereas deletion in MoMFs produced minimal metabolic effects. Single-cell transcriptomics revealed the molecular basis for this cell-type specificity - KCs exhibit a glucose-hungry metabolic phenotype that renders them uniquely dependent on Chi3l1-mediated regulation, while MoMFs maintain a relatively glucose-independent metabolic program. At the mechanistic level, we demonstrate that Chi3l1 functions as a physiological glucose sensor, directly binding extracellular glucose to limit its cellular uptake. This interaction establishes a crucial metabolic safeguard that specifically protects glucose-dependent KCs from lethal metabolic stress while sparing glucose-independent MoMFs. Through this precise modulation of glucose availability, Chi3l1 maintains metabolic homeostasis and preserves KCs populations during chronic dietary challenge (Figure 8).

**Figure 8.**
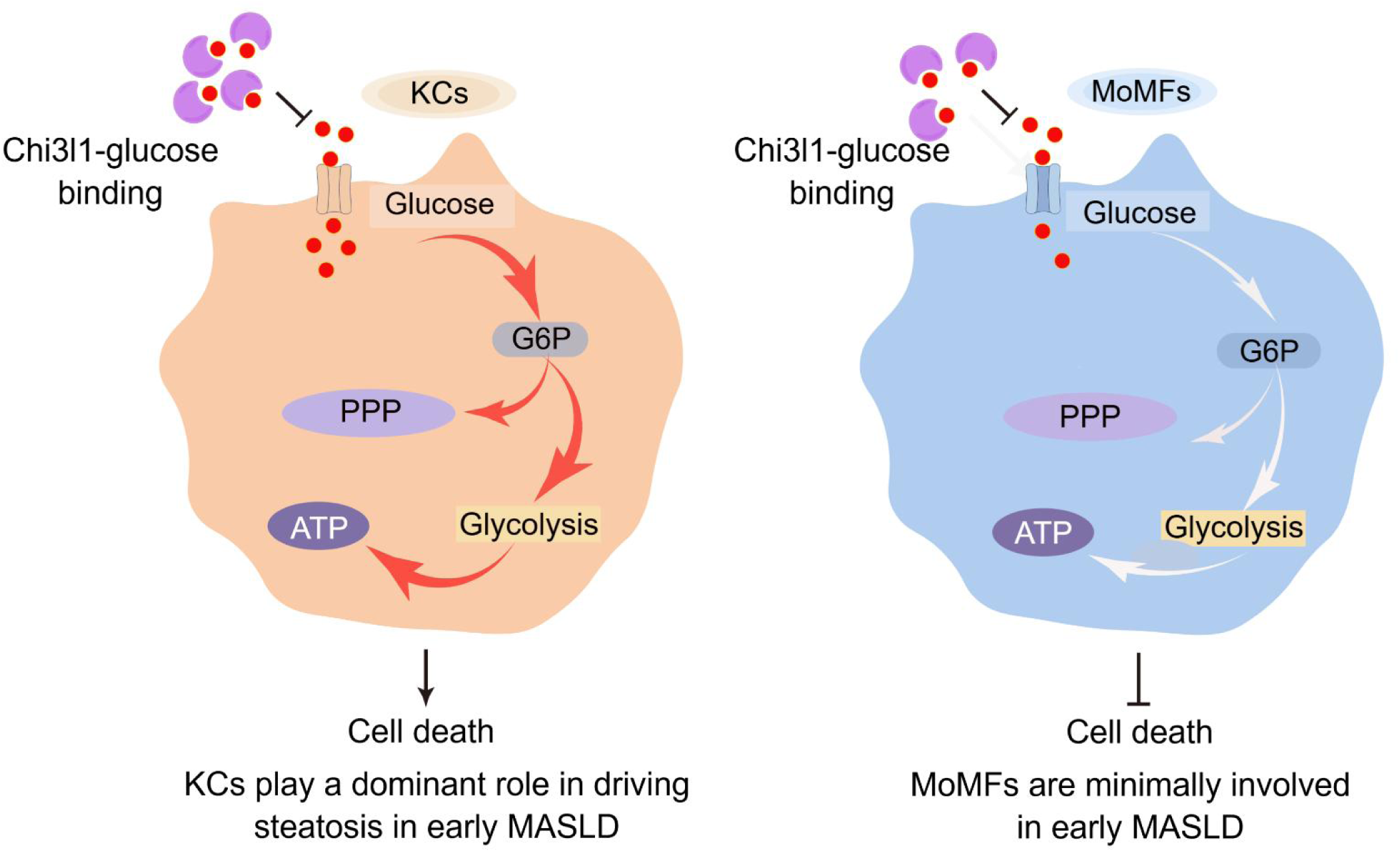
Differential regulation of KCs and MoMFs fate by Chi3l1-glucose interaction. KCs maintain a high-glucose activation state, while MoMFs exhibit a relatively low-glucose metabolic program. Chi3l1-glucose binding inhibits glucose uptake in KCs, thereby delaying KCs death and alleviating MASLD progression and metabolic dysfunction. In contrast, although Chi3l1-glucose binding similarly inhibits glucose uptake in MoMFs, their low basal glucose metabolism renders them resistant to this metabolic perturbation, resulting in minimal impact on MASLD pathogenesis.

Analysis of publicly available scRNA-seq datasets, including those from the Liver Atlas and prior studies^7–9^, indicates that Chil1 transcripts are mainly detected in neutrophils. In contrast, our immunofluorescence data show that Chi3l1 protein is predominantly localized in Kupffer cells under normal conditions and in both KCs and MoMFs during MASLD progression. This discrepancy likely reflects differences in transcript versus protein abundance and detection sensitivity. While scRNA-seq captures relative mRNA levels per cell, tissue-based staining reflects both expression and cell prevalence, highlighting macrophages as a major contributor to total hepatic Chi3l1 protein. Moreover, environmental factors such as diet, microbiota, or disease stage may influence Chil1 expression patterns across immune cell types.

Our study reveals fundamental differences in metabolic requirements between hepatic macrophage subsets that provide new insights into MASLD pathogenesis. We demonstrate that KCs and MoMFs play stage-specific roles in disease progression, with KCs serving as critical regulators of early metabolic homeostasis while MoMFs appear more involved in later inflammatory phases. This temporal specialization explains the striking dichotomy observed in our genetic models-KCs-specific Chi3l1 deletion dramatically exacerbated metabolic dysfunction, whereas MoMFs deletion showed minimal effects. The heightened glucose metabolism of KCs during MASLD renders them uniquely vulnerable to dietary stress. Chi3l1 serves as a crucial metabolic buffer in this context, directly protecting KCs through glucose modulation as evidenced by reduced glycogen accumulation and attenuated glycolytic flux. Our findings using the HFHC model complement previous findings in fibrogenic CDAA-HFAT models^27^ or MCD/CCL_4_ models^28^ or human livers^29^, collectively suggesting Chi3l1 may have dual roles in MASLD - maintaining metabolic balance through KCs in early disease while potentially influencing fibrogenesis via MoMFs in advanced stages. The accelerated KCs death in knockout models provides direct experimental evidence linking macrophage survival to metabolic outcomes, resolving key questions about MASLD progression mechanisms.

The structural characteristics of Chi3l1 have been extensively studied. Chi3l1 forms a homodimer, with each subunit containing a catalytic domain and a carbohydrate-binding domain. While the catalytic domain retains structural similarity to chitinases, it lacks enzymatic activity,^30,31^ and the carbohydrate-binding domain mediates interactions with carbohydrate ligands.^30^ While chitin-binding domains are traditionally known to interact with complex polysaccharides, our findings reveal that Chi3l1 (YKL-40), a mammalian chitinase-like protein, specifically binds to glucose—a simple monosaccharide. This represents a fundamental departure from canonical binding to insoluble polymers such as chitin and suggests a previously unrecognized role for Chi3l1 in monosaccharide recognition, potentially linking it to glucose metabolism and energy sensing. Furthermore, we observed that Chi3l1 protein levels increased in the serum of mice fed a high-fat, high-cholesterol (HFHC) diet for 16 weeks (Figure 6F) but plateaued with prolonged feeding (24 weeks; data not shown), suggesting an adaptive regulatory limit. Together, these findings indicate that Chi3l1 possesses glucose-binding capacity that may be functionally relevant but limited *in vivo*.

Our findings carry important translational potential for MASLD treatment. The discovery of Chi3l1’s glucose-sensing function in KCs suggests two complementary therapeutic strategies: first, developing Chi3l1-based interventions to preserve KC populations during early metabolic dysfunction; second, creating cell-type-specific approaches that selectively modulate glucose metabolism in KCs while sparing MoMFs. Importantly, although access to early-stage human liver tissue is limited due to the asymptomatic nature of the disease, multiple human studies have consistently reported elevated Chi3l1 levels in steatotic and fibrotic liver disease^28,32,33^, underscoring the clinical relevance of our mechanistic findings. Building on this evidence, the structural mapping of Chi3l1’s glucose-binding domain now enables rational design of small-molecule mimetics or biologics to therapeutically enhance this protective pathway. Besides, several key questions emerge for future research to advance these therapeutic possibilities: (1) How glucose levels are coordinated with other death inducers such as lipid toxicity; (2) Whether competing carbohydrate ligands modulate Chi3l1’s glucose-sensing capacity in different metabolic states; (3) Functional validation in primary human macrophages or human liver tissues would further strengthen the translational significance of this work. Addressing these questions will be crucial for translating our mechanistic insights into targeted therapies that account for the complex metabolic specialization of hepatic macrophage subsets.

Our findings reveal a novel metabolic checkpoint in which Chi3l1 selectively sustains KCs populations by modulating glucose metabolism, offering key insights into MASLD pathogenesis. The study highlights the therapeutic potential of targeting Chi3l1-glucose interactions to preserve protective KCs and curb MASLD progression. Future research should explore whether Chi3l1 supplementation or pharmacological modulation can rescue KCs viability, as well as investigate whether this mechanism extends to other macrophage-driven metabolic disorders, such as MASH or diabetes. By identifying cell type-specific metabolic vulnerabilities, this work paves the way for precision therapies that selectively manipulate macrophage subsets to treat liver disease.

## Abbreviations

MASLD: metabolic dysfunction-associated steatotic liver disease
MASH: metabolic dysfunction-associated steatohepatitis
KCs: Kupffer cells
Chi3l1: Chitinase 3 like 1
ERK1/2: extracellular signal-regulated kinase ½
PI3K: phosphoinositide-3 kinase
ALT: Alanine aminotransferase
AST: aspartate aminotransferase
TC: cholesterol
TG: triglyceride
NPCs: nonparenchymal cells
HFHC: high fat high cholesterol diet
NCD: normal chow diet
Clec4f: C-type lectin domain family 4
TIM4: T cell immunoglobulin mucin protein 4
MoMFs: monocyte-derived macrophages
HFD: high-fat diet
MCD: methionine/choline deficient diet
WD: western diet
PPP: pentose phosphate pathway
BMDM: bone marrow derived macrophages
DMSO: dimethyl sulfoxide
MAFL: non-alcoholic fatty liver
rChi3l1: recombinant murine Chi3l1
PA: palmic acid
Iso: Isopropyl alcohol
IGTT: intraperitoneal glucose tolerance test
ITT: insulin tolerance test
scRNA-seq: single-cell RNA sequencing
MoKCs: monocytes-derived Kupffer cells
ALD: alcohol-induced liver disease
AILI: acetaminophen-induced liver injury
EmKCs: embryo-derived Kupffer cells
DT: diphtheria toxin
WT: wild-type
TUNEL: TdT-mediated dUTP Nick-End Labeling

## Acknowledgements

We thank Dr. Bin Qi (Yunnan University) for suggestions and discussion. We thank Guangxun Meng (The Shanghai Institute of Immunity and Infection of the Chinese Academy of Sciences) for providing us with L929 cells. We thank Cynthia Ju (UTHealth) for advice in manuscript submission.

## Author Contributions

JH conducted the experiments, analyzed the data, and wrote the manuscript. BC performed the scRNA seq analysis and wrote the manuscript. WJL performed scRNA seq analysis during the revision. WX helped mice care and feeding. RXY helped scRNA-seq library preparation. CXD conducted the initial analysis of the scRNA-seq data under the supervision of CP. XEZ participated in sample collection. KQW and LW purified the recombinant Chi3l1 protein. RZY drew molecular models of glucose and chitin. CX and RL helped with 2-NBDG *in vivo* imaging. CPL and XKL helped synthesize biotin-labeled glucose. ZS conceived, organized, and designed the study, and wrote the manuscript.

## Main figure

**Figure S1.**
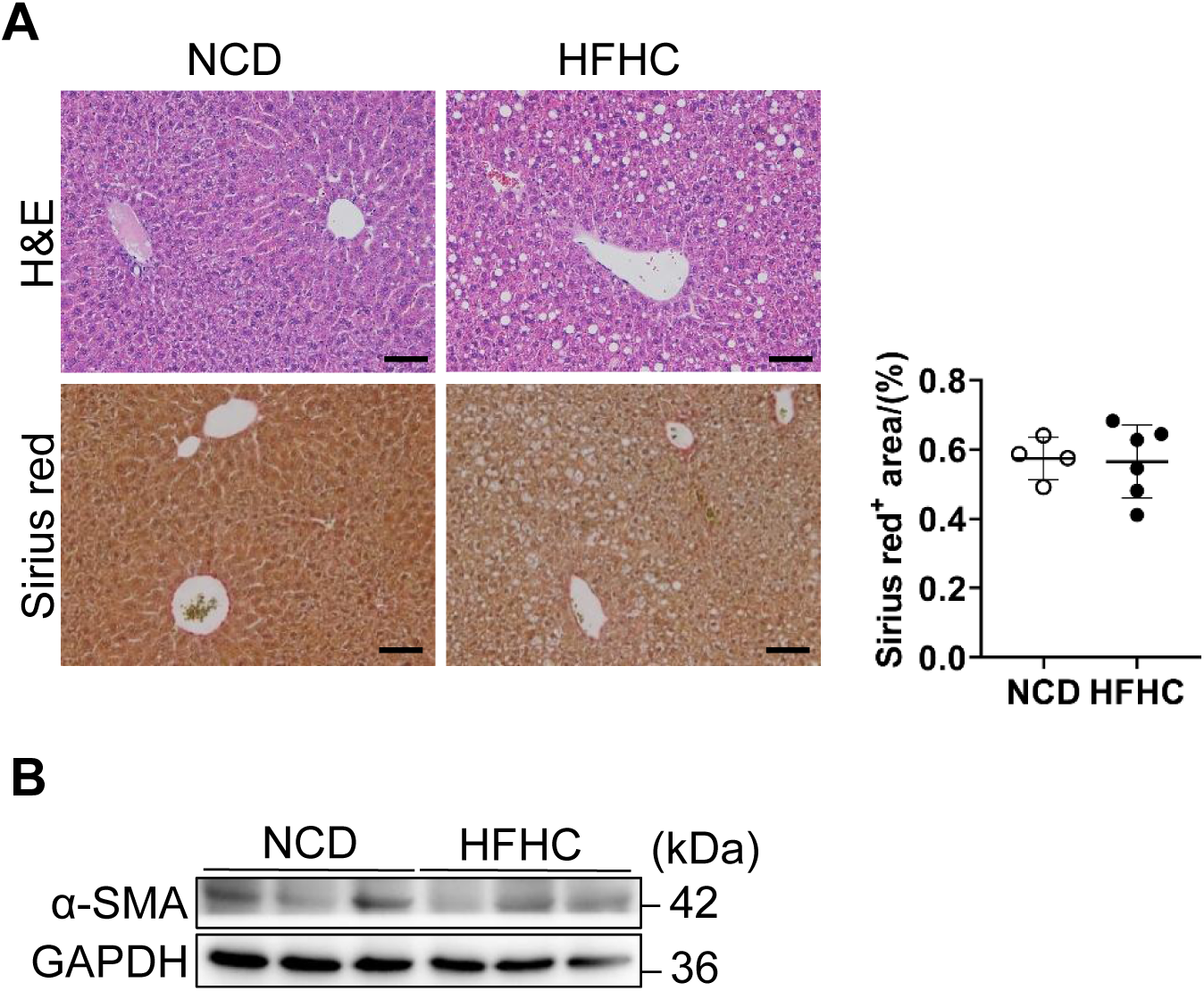
MASLD progression in the HFHC diet-induced mouse model. **(A)** Representative liver sections from wild-type C57BL/6J mice fed either a normal chow diet (NCD) or a high-fat, high-cholesterol (HFHC) diet for 16 weeks. H&E and Sirius Red staining were used to assess lipid deposition, inflammation and fibrosis. Scale bar: 20 µm. Quantification of Sirius Red–positive area is shown. **(B)** Western blot analysis of α-SMA expression in whole liver lysates from NCD-and HFHC-fed mice (n = 3 mice/group) to evaluate activation of hepatic stellate cells.

**Figure S2.**
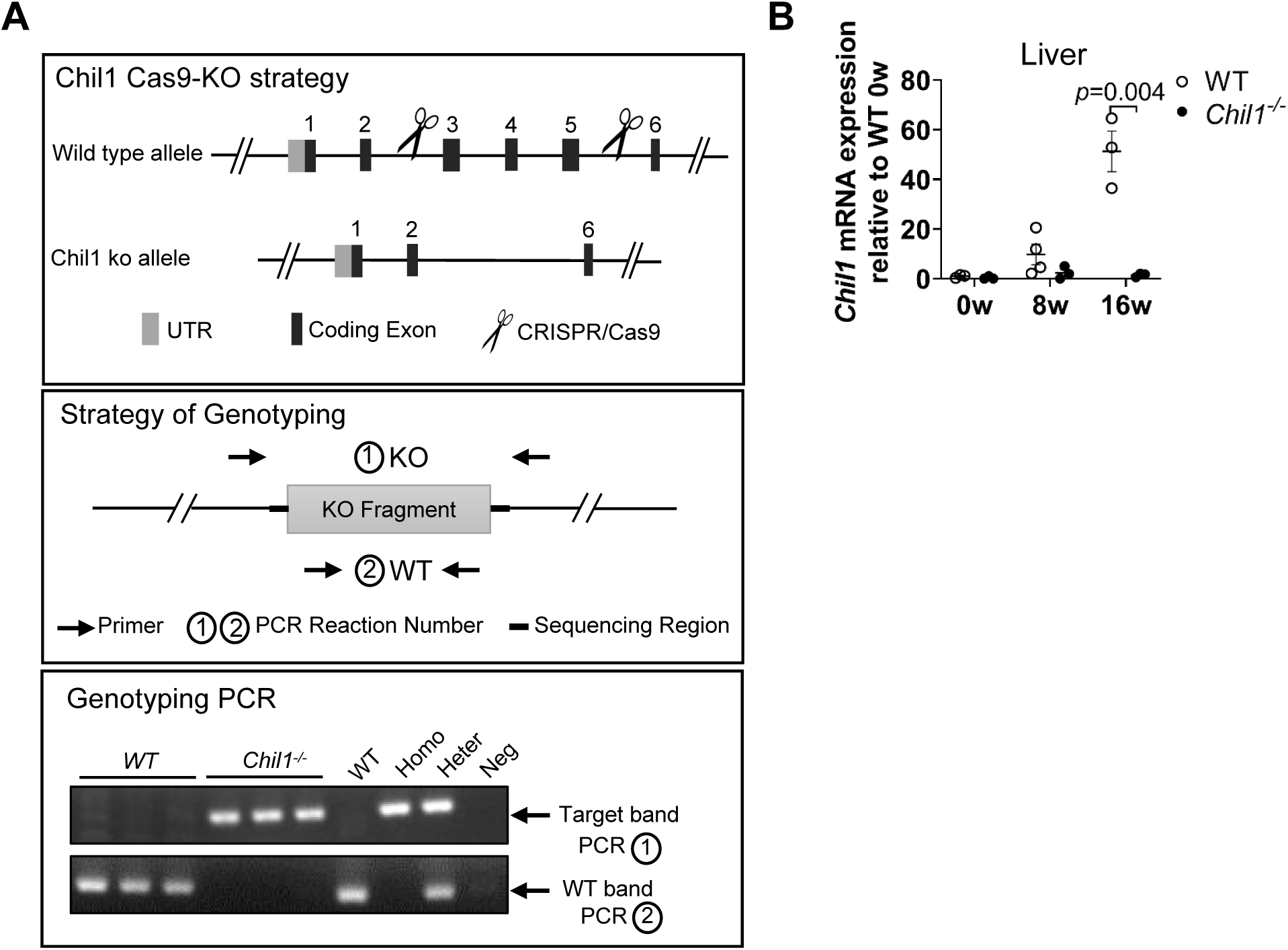
Generation and validation of *Chil1^-/-^* mice. **(A)** The construction, genotyping strategy and genotyping results of *Chil1^-/-^* mice. P: positive control; WT: Wild-type; Neg: Blank control(ddH_2_O). **(B)** qRT-PCR analysis of mRNA expression levels of Chil1 in liver tissues of WT and *Chil1^-/-^* mice fed with HFHC for 0, 8 and 16 weeks. n=3-4 mice/group. Two-tailed, unpaired student t-test was performed in B. P value is as indicated.

**Figure S3.**
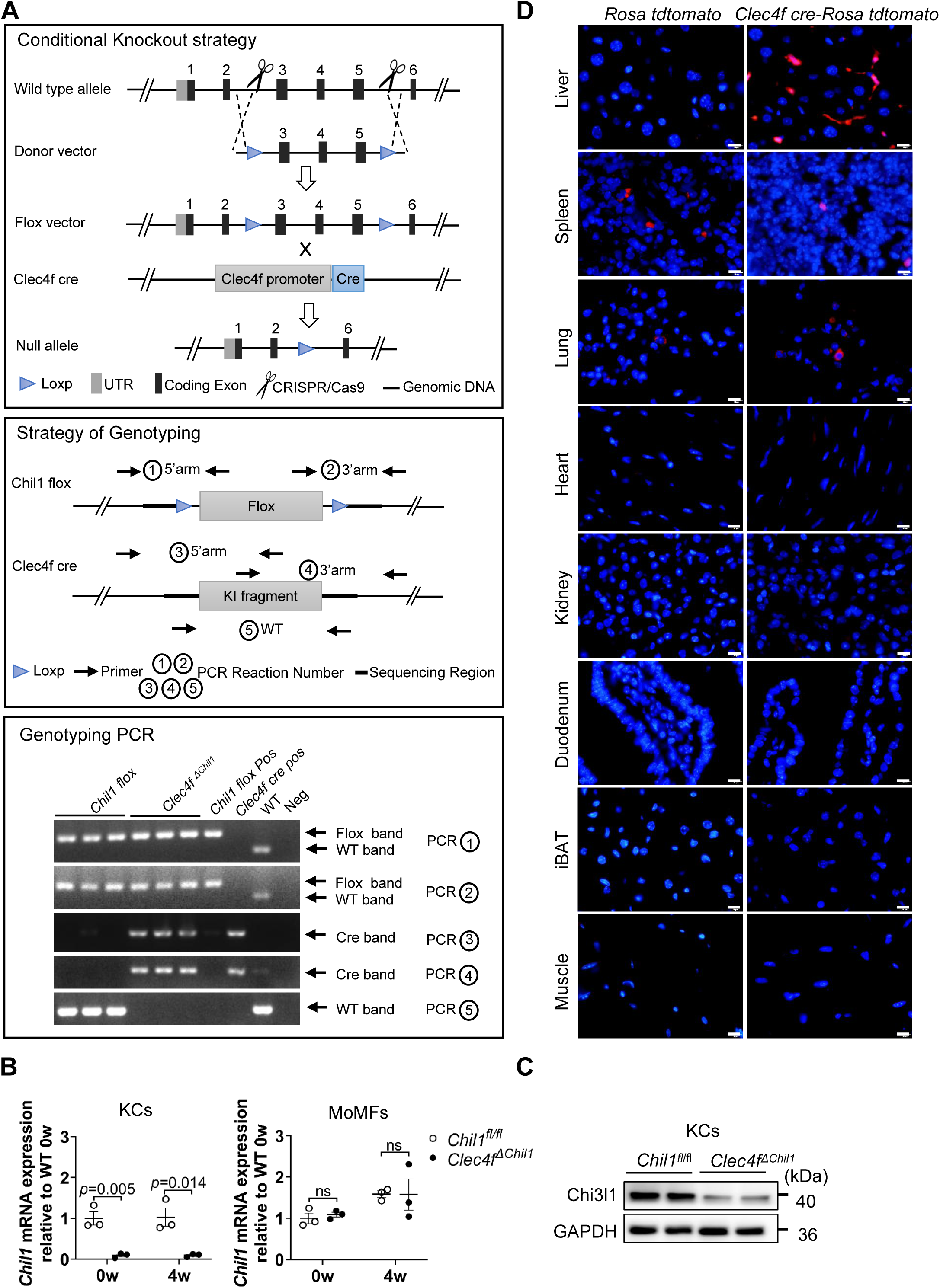
The construction and genotype of *Clec4f^ΔChil^*^1^ mice. **(A)** The construction, genotyping strategy and genotyping results of *Clec4f^ΔChil^*^1^ mice. P: positive control; WT: Wild-type; Neg: Blank control (ddH_2_O). **(B)** qRT-PCR analysis of mRNA expression levels of *Chil1* in KCs (CD45^+^ F4/80^hi^ CD11b^low^ TIM4^hi^) or MoMFs (CD45^+^ F4/80^low^ CD11b^hi^ Ly6G^-^ TIM4^-^) FACS sorted from *Chil1^fl/fl^* and *Clec4f^ΔChil^*^1^ mice at 0 and 4 weeks post HFHC diet. n=3 mice/group. **(C)** Western blot to detect Chi3l1 expression in isolated KCs of *Chil1^fl/fl^* and *Clec4f^ΔChil^*^1^ mice. n=2 mice/group. **(D)** The expression specificity of Clec4f was examined in various tissues in *Clec4f cre-Rosa tdtomato* mice, which is generated by crossing *Clec4f-cre* with *Rosa-tdtomato* mice.

**Figure S4.**
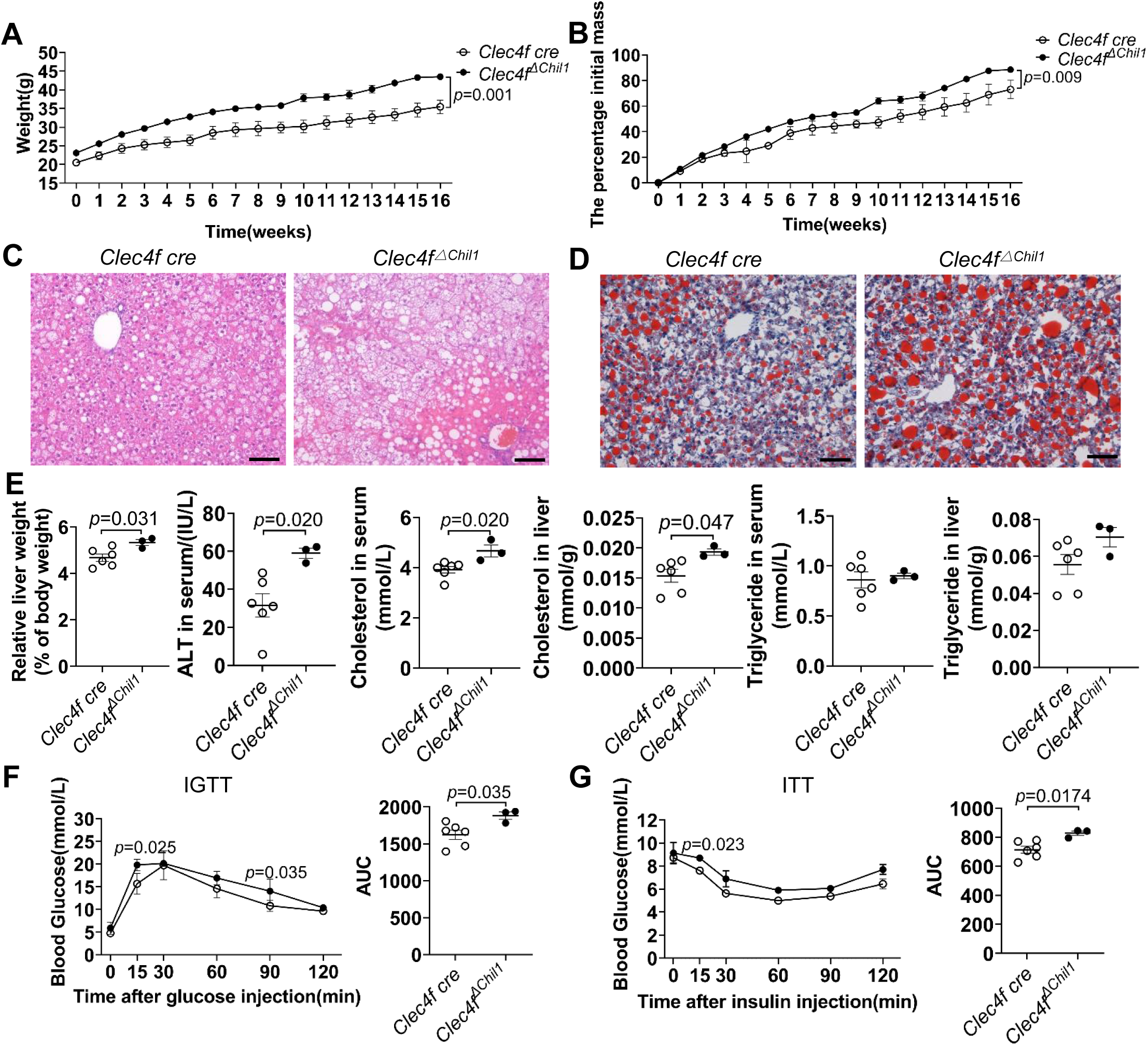
Deficiency of Chi3l1 in Kupffer cells promotes insulin resistance and hepatic lipid accumulation. *Clec4f cre* and *Clec4f^ΔChil^*^1^ mice were fed with a HFHC diet for 16 weeks. **(A, B)** Body weight was recorded during HFHC diet feeding (A) and expressed as a percentage of initial body mass (B). **(C, D)** H&E (C) and oil red o staining (D) was performed to examine liver histology and hepatic lipid accumulation in in both genotypes after 16 weeks of HFHC diet. Scale bar = 20 µm. **(E)** Liver index (liver weight/body weight × 100%), ALT levels and serum and liver Cholesterol or Triglyceride levels were measured in both genotypes after 16 weeks of HFHC diet. n=3-6 mice/group. **(F&G)** Intraperitoneal glucose tolerance test (IGTT) and insulin tolerance test (ITT) were performed after 16 weeks of HFHC feeding in both genotypes.n=3-6 mice/group. Representative images were shown in C, D. Two-tailed, unpaired student t-test was performed in A,B,E-G. P value is as indicated.

**Figure S5.**
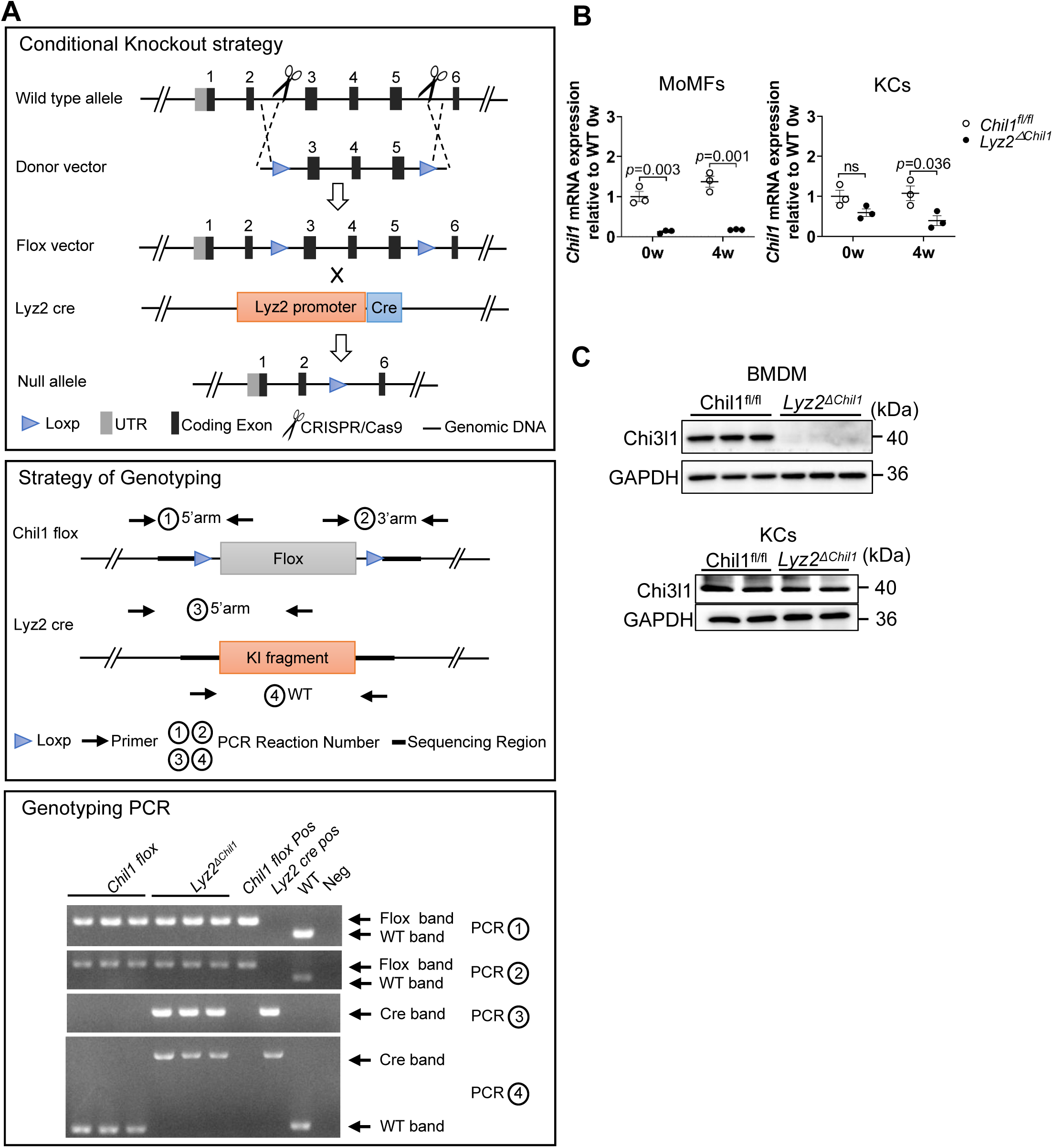
The construction and genotype of *Lyz2^ΔChil^*^1^ mice. **(A)** The construction, genotyping strategy and genotyping results of *Lyz2^ΔChil^* ^1^ mice. pos: positive control; WT: Wild-type; Neg: Blank control(ddH2O). **(B)** qRT-PCR analysis of mRNA expression levels of *Chil1* in KCs (CD45^+^ F4/80^hi^ CD11b^low^ TIM4^hi^) or MoMFs (CD45^+^ F4/80^low^ CD11b^hi^ Ly6G^-^ TIM4^-^) FACS sorted from *Chil1^fl/fl^* and *Lyz2^ΔChil^*^1^ mice at 0 and 4 weeks post HFHC diet. n= 3 mice/group. **(C)** Western blotting analysis of protein levels of Chi3l1 in BMDM and primary KCs of *Chil1^fl/f l^* and *Lyz2^ΔChil^*^1^ mice. n=2-3 mice/group.

**Figure S6.**
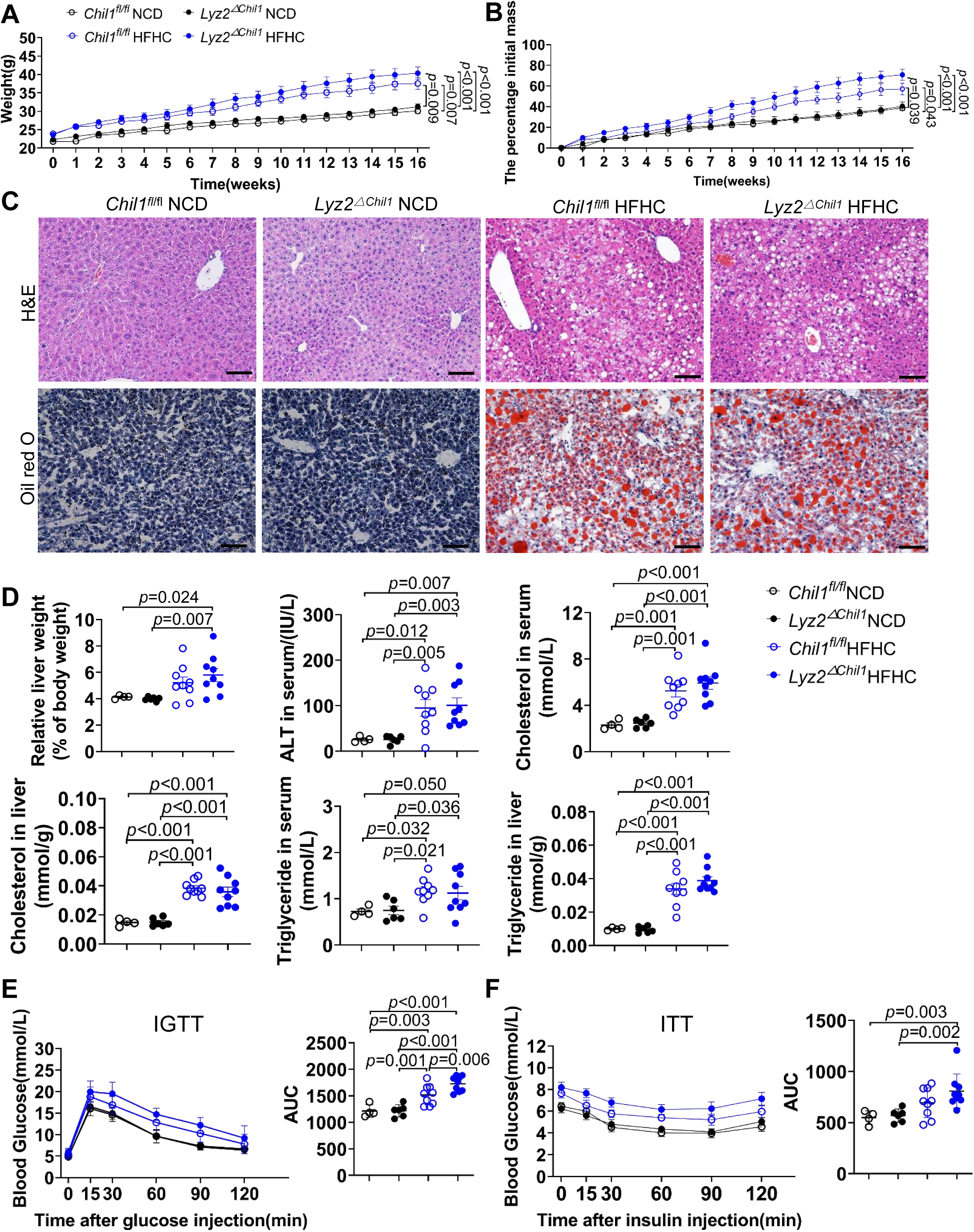
Deficiency of Chi3l1 in MoMFs barely affect insulin resistance and hepatic lipid accumulation. *Chil1^fl/fl^* and *Lyz2^ΔChil^*^1^ mice were fed either a normal chow diet (NCD) or a high-fat, high-cholesterol (HFHC) diet for 16 weeks. **(A, B)** Body weight was recorded during HFHC diet feeding (A) and expressed as a percentage of initial body mass (B). **(C)** H&E (Upper panel) and oil red o staining (Lower panel) was performed to examine liver histology and hepatic lipid accumulation in both genotypes after 16 weeks of NCD or HFHC diet. Scale bar = 20 µm. **(D)** Liver index (liver weight/body weight × 100%), ALT levels, and serum and liver Cholesterol or Triglyceride levels were measured in both genotypes after 16 weeks on NCD or HFHC diets. n=4-9 mice/group. **(E, F)** Intraperitoneal glucose tolerance test (IGTT) and insulin tolerance test (ITT) were performed after 16 weeks of NCD or HFHC feeding in both genotypes (n = 4–9 mice per group). Representative images were shown in (C). One-way ANOVA was performed in (A, B, D-F). P-values are as indicated.

**Figure S7.**
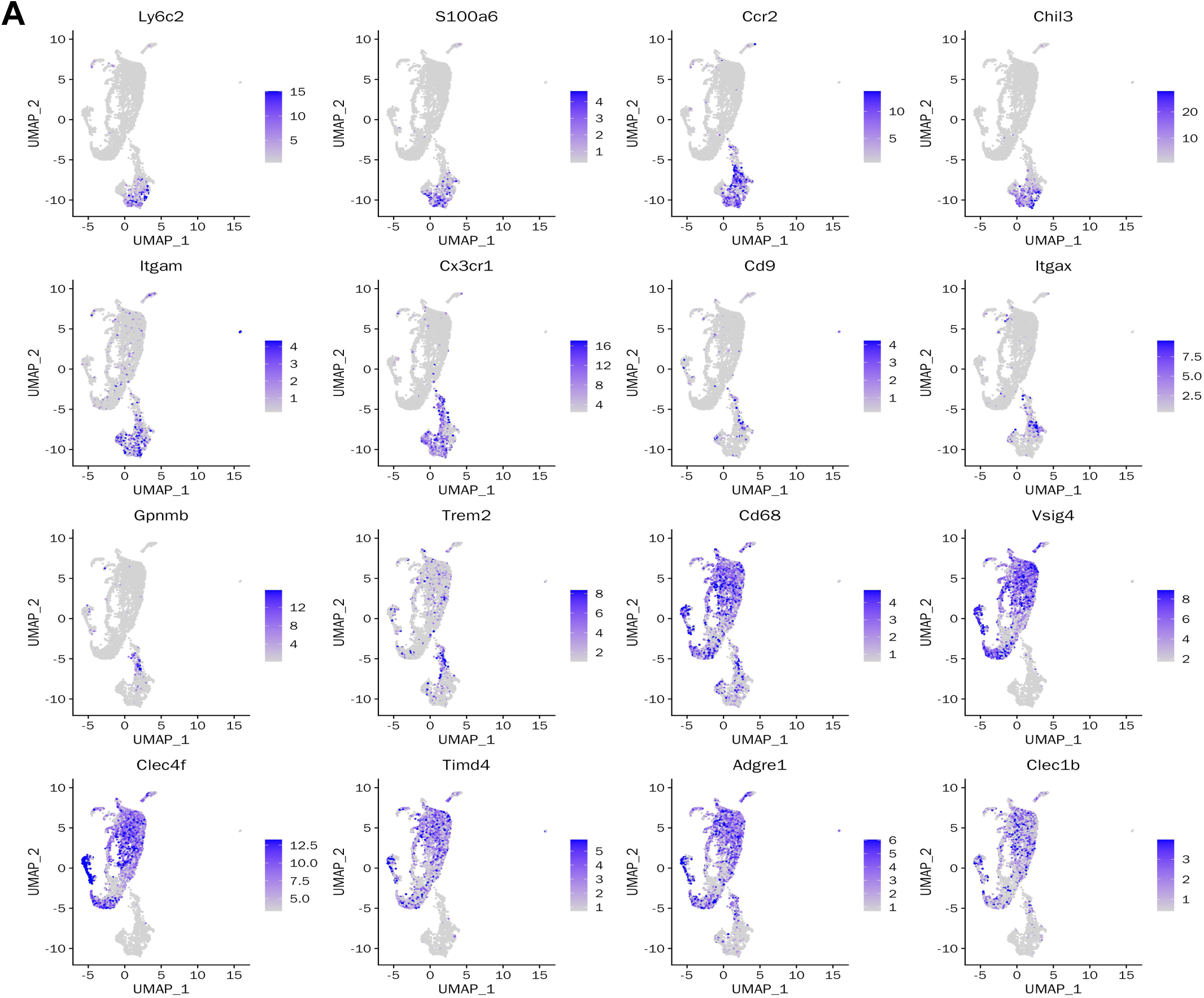
Gene expression levels of lineage-specific marker genes in monocytes/macrophages clusters. **(A)** Scaled gene expression levels of each lineage-specific marker gene are shown in UMAP plots of monocytes/macrophages clusters. Colors indicate gene expression levels.

**Figure S8.**
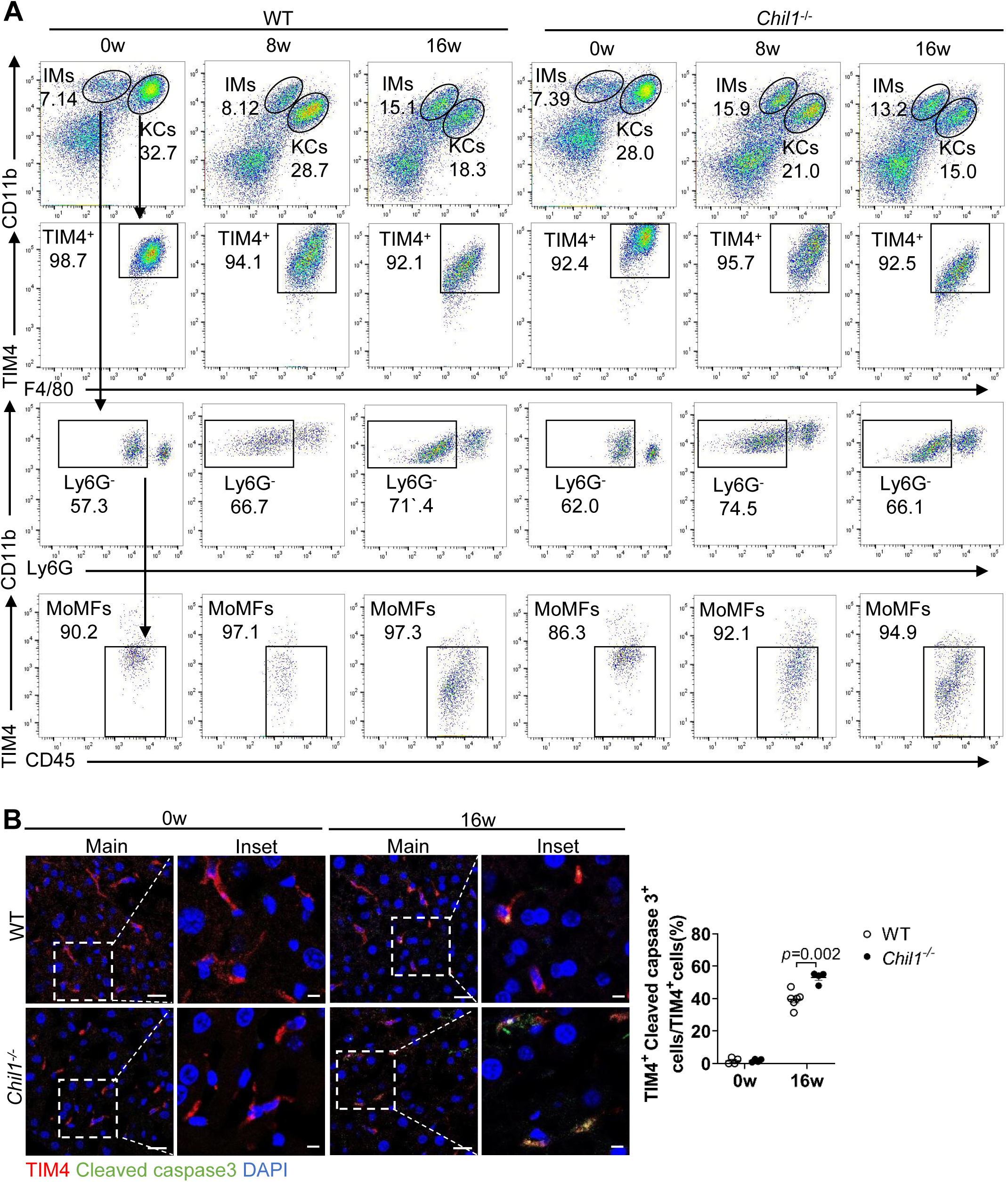
Chi3l1 deficiency promote KCs death during MASLD. WT and *Chil1^-/-^* mice were fed with a HFHC diet for 0, 8, 16 weeks. **(A)** Flow cytometry analysis of KCs (CD45^+^ F4/80^hi^ CD11b^low^ TIM4^hi^) and MoMFs (CD45^+^ F4/80^low^ CD11b^hi^ Ly6G^-^ TIM4^-^) among NPCs between WT and *Chil1^-/-^* mice. **(B)** Immunofluorescent staining to detect TIM4(red), Cleaved caspase3(green), and nuclear DAPI (blue) in liver sections. Scale bar=20μm and 5μm(Insets). Cleaved caspase3^+^ TIM4^+^ cells/ TIM4^+^ cells were statistically analyzed. n=4-6 mice/group. Representative images are shown in A, B. Student t-test was performed in B. P value is as indicated.

**Figure S9.**
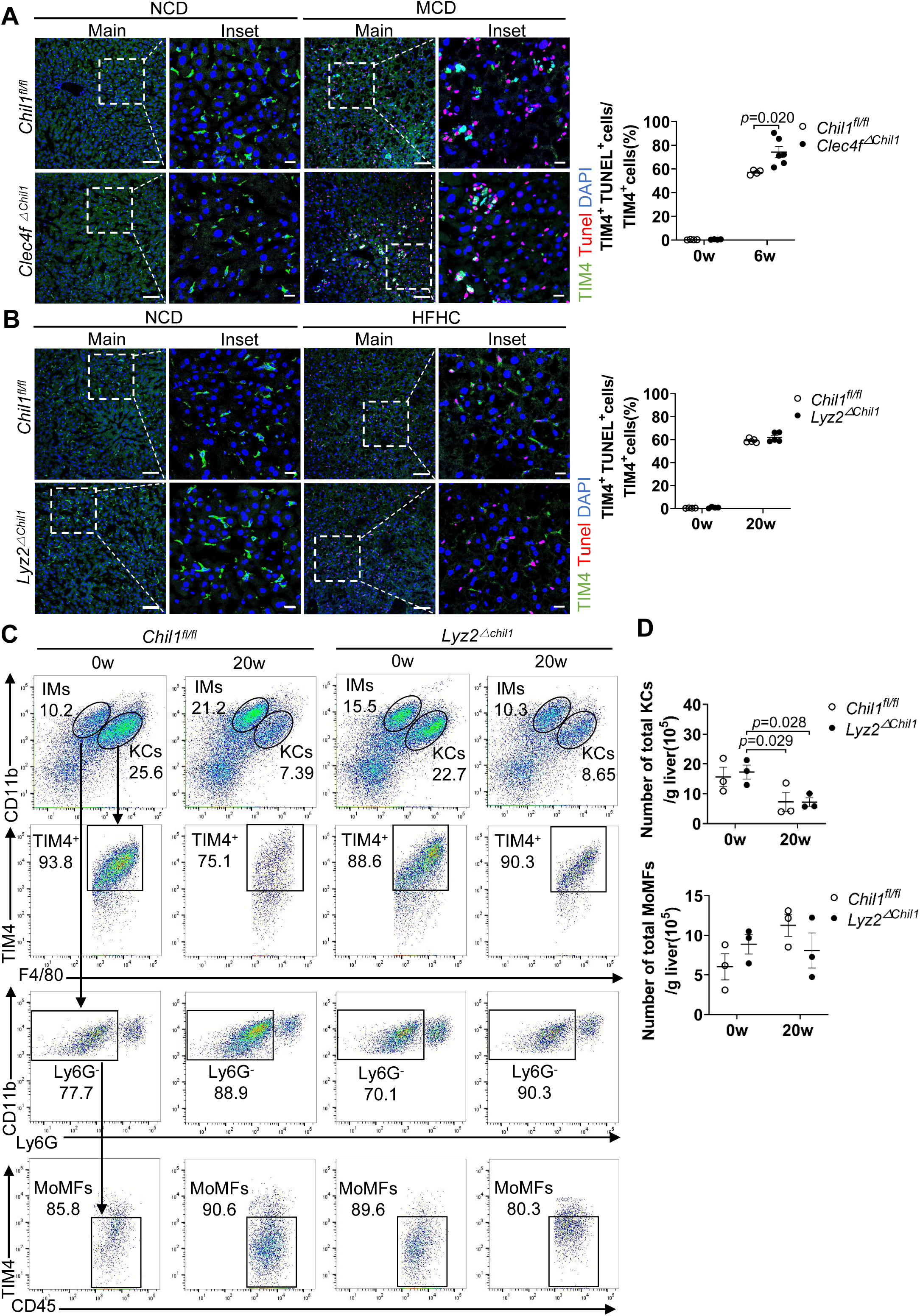
Deficiency of Chi3l1 in KCs but not MoMFs promote KCs death during MASLD. *Chil1^fl/fl^* and *Clec4f^ΔChil^*^1^ mice were fed with a MCD diet for 6 weeks. *Chil1^fl/fl^* and *Lyz2^ΔChil^*^1^ mice were fed with a HFHC diet for 20 weeks. **(A)** Immunofluorescent staining to detect TIM4(green), TUNEL(red), and nuclear DAPI (blue) in liver sections of Chil1^fl*/fl*^ and *Clec4f^ΔChil^*^1^ mice. Scale bar=50µm and 20µm (Insets). TUNEL^+^ TIM4^+^ cells/TIM4^+^ cells were statistically analyzed. n=4-6 mice/group. **(B)** Immunofluorescent staining to detect TIM4(green), TUNEL(red), and nuclear DAPI (blue) in liver sections of *Chil1^fl/fl^* and *Lyz2^ΔChil^*^1^ mice. Scale bar=50µm and 20µm (Insets). TUNEL^+^ TIM4^+^ cells/TIM4^+^ cells were statistically analyzed. n=4-5 mice/group. **(C)** Flow cytometry analysis of KCs (CD45^+^ F4/80^hi^ CD11b^low^ TIM4^hi^) and MoMFs (CD45^+^ F4/80^low^ CD11b^hi^ Ly6G^-^ TIM4^-^) among NPCs between *Chil1^fl/fl^* and *Lyz2^ΔChil^*^1^ mice. **(D)** Number of KCs or MoMFs/gram(g) liver were statistically analyzed. n= 3 mice/group. Representative images are shown in A-C. Student t-test was performed in A-B. One-way ANOVA was performed in D. P value is as indicated.

## Reference

1 H Hardy T, Oakley F, Anstee QM, et al. Nonalcoholic Fatty Liver Disease: Pathogenesis and Disease Spectrum. Annu Rev Pathol-Mech 11, 451–496, doi:10.1146/annurev-pathol-012615-044224 (2016).

2 Eslam M, Newsome PN, Sarin SK, et al. A new definition for metabolic dysfunction-associated fatty liver disease: An international expert consensus statement. Journal of Hepatology 73, 202–209, doi:10.1016/j.jhep.2020.03.039 (2020).

3 Kazankov K, Jørgensen SMD, Thomsen KL, et al. The role of macrophages in nonalcoholic fatty liver disease and nonalcoholic steatohepatitis. Nature reviews. Gastroenterology & hepatology 16, 145–159, doi:10.1038/s41575-018-0082-x (2019).

4 Gomez Perdiguero E, Klapproth K, Schulz C, et al. Tissue-resident macrophages originate from yolk-sac-derived erythro-myeloid progenitors. Nature 518, 547–551, doi:10.1038/nature13989 (2015).

5 Hashimoto D, Chow A, Noizat C, et al. Tissue-Resident Macrophages Self-Maintain Locally throughout Adult Life with Minimal Contribution from Circulating Monocytes. Immunity 38, 792–804, doi:10.1016/j.immuni.2013.04.004 (2013).

6 Tran S, Baba I, Poupel L, et al. Impaired Kupffer Cell Self-Renewal Alters the Liver Response to Lipid Overload during Non-alcoholic Steatohepatitis. Immunity 53, 627–640.e625, doi:10.1016/j.immuni.2020.06.003 (2020).

7 Daemen S, Gainullina A, Kalugotla G, et al. Dynamic Shifts in the Composition of Resident and Recruited Macrophages Influence Tissue Remodeling in NASH. Cell Rep 34, 108626, doi:10.1016/j.celrep.2020.108626 (2021).

8 Seidman JS, Troutman TD, Sakai M, et al. Niche-Specific Reprogramming of Epigenetic Landscapes Drives Myeloid Cell Diversity in Nonalcoholic Steatohepatitis. Immunity 52, 1057, doi:10.1016/j.immuni.2020.04.001 (2020).

9 Remmerie A, Martens L, Thoné T, et al. Osteopontin Expression Identifies a Subset of Recruited Macrophages Distinct from Kupffer Cells in the Fatty Liver. Immunity 53, 641–657.e614, doi:10.1016/j.immuni.2020.08.004 (2020).

10 Green DR, Galluzzi L, Kroemer G. Cell biology. Metabolic control of cell death. Science 345, 1250256, doi:10.1126/science.1250256 (2014).

11 Inomata Y, Oh JW, Taniguchi K, et al. Downregulation of miR-122-5p Activates Glycolysis via PKM2 in Kupffer Cells of Rat and Mouse Models of Non-Alcoholic Steatohepatitis. International journal of molecular sciences 23, doi:10.3390/ijms23095230 (2022).

12 Lodge M, Scheidemantle G, Adams VR, et al. Fructose regulates the pentose phosphate pathway and induces an inflammatory and resolution phenotype in Kupffer cells. Sci Rep-Uk 14, doi:ARTN 402010.1038/s41598-024-54272-w (2024).

13 Dong T, Hu GG, Fan ZQ, et al. Activation of GPR3-β-arrestin2-PKM2 pathway in Kupffer cells stimulates glycolysis and inhibits obesity and liver pathogenesis. Nature Communications 15, doi:ARTN 80710.1038/s41467-024-45167-5 (2024).

14 Dong T, Hu GG, Fan ZQ, et al. Role of chitin and chitinase/chitinase-like proteins in inflammation, tissue remodeling, and injury. Annual review of physiology 73, 479–501, doi:10.1146/annurev-physiol-012110-142250 (2011).

15 Dela Cruz CS, Liu W, He CH, et al. Chitinase 3-like-1 promotes Streptococcus pneumoniae killing and augments host tolerance to lung antibacterial responses. Cell host & microbe 12, 34–46, doi:10.1016/j.chom.2012.05.017 (2012).

16 He CH, Lee CG, Dela Cruz CS, et al. Chitinase 3-like 1 regulates cellular and tissue responses via IL-13 receptor α2. Cell reports 4, 830–841, doi:10.1016/j.celrep.2013.07.032 (2013).

17 Zhou Y, He CH, Herzog EL, et al. Chitinase 3-like-1 and its receptors in Hermansky-Pudlak syndrome-associated lung disease. The Journal of clinical investigation 125, 3178–3192, doi:10.1172/jci79792 (2015).

18 Lee CG, Hartl D, Lee GR, et al. Role of breast regression protein 39 (BRP-39)/chitinase 3-like-1 in Th2 and IL-13-induced tissue responses and apoptosis. J Exp Med 206, 1149–1166, doi:10.1084/jem.20081271 (2009).

19 Fu JT, Liu J, Wu WB, et al. Targeting EFHD2 inhibits interferon-γ signaling and ameliorates non-alcoholic steatohepatitis. Journal of Hepatology 81, doi:10.1016/j.jhep.2024.04.009 (2024).

20 Jeelani I, Moon JS, da Cunha FF, et al. HIF-2α drives hepatic Kupffer cell death and proinflammatory recruited macrophage activation in nonalcoholic steatohepatitis. Science Translational Medicine 16, doi:ARTN eadi028410.1126/scitranslmed.adi0284 (2024).

21 Shan Z, Li L, Atkins CL, et al. Chitinase 3-like-1 contributes to acetaminophen-induced liver injury by promoting hepatic platelet recruitment. Elife 10, doi:10.7554/eLife.68571 (2021).

22 Scott CL, T’Jonck W, Martens L, et al. The Transcription Factor ZEB2 Is Required to Maintain the Tissue-Specific Identities of Macrophages. Immunity 49, 312–325 e315, doi:10.1016/j.immuni.2018.07.004 (2018).

23 Li L, Cui L, Lin P, et al. Kupffer-cell-derived IL-6 is repurposed for hepatocyte dedifferentiation via activating progenitor genes from injury-specific enhancers. Cell Stem Cell 30, 283-+, doi:10.1016/j.stem.2023.01.009 (2023).

24 Green DR, Galluzzi L, Kroemer G. Metabolic control of cell death. Science 345, 1466, doi:ARTN 125025610.1126/science.1250256 (2014).

25. 25 He J, Li R, Xie Cheng, et al. Hyperactivated Glycolysis Drives Spatially-Patterned Kupffer Cell Depletion in MASLD. bioRxiv, doi:doi: 10.1101/2025.09.26.678483 (2025).

26 Liu QX, Li JX, Zhang WJ, et al. Glycogen accumulation and phase separation drives liver tumor initiation. Cell 184, 5559, doi:10.1016/j.cell.2021.10.001 (2021).

27 Kim AD, Kui L, Kaufmann B, et al. Myeloid-specific deletion of chitinase-3-like 1 protein ameliorates murine diet-induced steatohepatitis progression. J Mol Med 101, 813–828, doi:10.1007/s00109-023-02325-4 (2023).

28 Higashiyama M, Tomita K, Sugihara N, et al. Chitinase 3-like 1 deficiency ameliorates liver fibrosis by promoting hepatic macrophage apoptosis. Hepatology Research 49, 1316–1328, doi:10.1111/hepr.13396 (2019).

29 Nishimura N, De Battista D, McGivern DR, et al. Chitinase 3-like 1 is a profibrogenic factor overexpressed in the aging liver and in patients with liver cirrhosis. P Natl Acad Sci USA 118, doi:ARTN e201963311810.1073/pnas.2019633118 (2021).

30 Fusetti F, Pijning T, Kalk KH, et al. Crystal structure and carbohydrate-binding properties of the human cartilage glycoprotein-39. Journal of Biological Chemistry 278, 37753–37760, doi:10.1074/jbc.M303137200 (2003).

31 Houston DR, Recklies AD, Krupa JC, et al. Structure and ligand-induced conformational change of the 39-kDa glycoprotein from human articular chondrocytes. Journal of Biological Chemistry 278, 30206–30212, doi:10.1074/jbc.M303371200 (2003).

32 Huang X, Zhuang J, Yang Y, et al. Diagnostic Value of Serum Chitinase-3-Like Protein 1 for Liver Fibrosis: A Meta-analysis. Biomed Res Int 2022, 3227957, doi:10.1155/2022/3227957 (2022).

33 Liu S, Peng C, Xia S, et al. Chitinase 3-like protein 1: a diagnostic biomarker for early liver fibrosis in autoimmune liver diseases. Front Immunol 16, 1504066, doi:10.3389/fimmu.2025.1504066 (2025).

