## Supplementary materials and methods for "Differential Regulation of Hepatic Macrophage Fate by Chi3l1 in MASLD"

### Animal experiments and procedures

*Intraperitoneal glucose tolerance test (IGTT)&Intraperitoneal insulin tolerance test (ITT)* For the IGTT, the mice were fasted for approximately 13-14 hours while having access to water. They were then intraperitoneally injected with saline containing 20% glucose (1g/kg of body weight, Cat# 10057153461926). For the ITT, the mice were fasted for approximately 5 h with access to water. They were then intraperitoneally injected with insulin (0.1 U/mL, 0.75 U per kg of body weight, Cat# 10090506507183). During both tests, the blood glucose levels were measured at 0, 30, 60, 90, and 120 min using a blood glucose meter (Roche ACCU-CHEK®Performa). Care was taken to collect blood from the tip of the tail by using gentle movements to avoid stress-induced hyperglycemia. After the tests, the mice were placed in a clean cage with food and water available and monitored for 1 hour. To ensure accurate and reliable results, we conducted any subsequent ITTs at least one week after the previous test to avoid any interference.

*Genotyping* Sample preparation and procedure were conducted as previously described.[1] Primer sequences are provided in Table S1.

**Table S1**

| Aim genes | PCR No. | Primer No. | Sequence | Band Size |
|---|---|---|---|---|
| *Chil1$^{flox//flox}$* | ① 5'arm | F1(JS05000-Chil1-5wt-tF1) | CTTGTTCAGCCAAGGTGATGGGTA | WT:270bp targeted:375bp |
| | | R1(JS05000-Chil1-5wt-tR1) | CTACCTGATTGCTGGGGCTCATTA | |
| | ② 3'arm | F2(JS15000-Chil1-3wt-F1) | CCAGTATTTAGAGGCAGAGAGATGGTG | WT:283bp targeted:384bp |
| | | R2(JS15000-Chil1-3wt-R1) | CTCGAATTCAGAAATCTGCCTGCCT | |
| *Chil1$^{-/-}$* | ① | F1(JS05000-Chil1-5wt-tF1) | CTTGTTCAGCCAAGGTGATGGGTA | WT: 2960 bp Targeted: ~256 bp |
| | | R1(JS05000-Chil1-3wt-tR1) | CTCGAATTCAGAAATCTGCCTGCCT | |
| | ② | F2(JS15000-Chil1-wt-F1) | CTGTTAGTTGCACCTTGGAGCAGTCA | WT: 309 bp Targeted:0 bp |
| | | R2(JS15000-Chil1-wt-R1) | CAGATATAGGAGAACATCCAGTCTGGG | |
| *Clec4f Cre* | ①5'arm | GPS00003712-Clec4f-wt-tF1 | CCCATCCTGAGGTCTCTTTATGC | WT:0bp Targeted:258bp |
| | | IRES-tR2 | TAGAGTCCAGATCTTCCGGGTAC | |
| | ②3'arm | iCre-tF1 | GGCTGGACCAATGTGAACATTG | WT:0bp Targeted:265bp |
| | | GPS00003712-Clec4f-wt-tR1 | TATTGAGGGCTTATCTGGGCAG | |
| | ③WT | GPS00003712-Clec4f-wt-tF1A | GGAGAGCGAGAAGACTGTGTTCAC | WT:250bp Targeted:1927bp |
| | | GPS00003712-Clec4f-wt-tR1A | GACTCCAATGCAGGGCTTGTCT | |

| | | | | |
|---|---|---|---|---|
| *Lyz2 Cre* | ①5'arm | F1 | AGTGCTGAAGTCCATAGATCGG | WT:0bp Targeted:258bp |
| | | R1 | CTGATTCTCCTCATCACCAGG | |
| | ②WT | F2 | AGTGCTGAAGTCCATAGATCGG | WT:0bp Targeted:265bp |
| | | R2 | GTCACTCACTGCTCCCCTGT | |

**Isolation of NPCs and KCs**

Hepatic nonparenchymal cells (NPCs) were isolated as previously described.[2] In brief, the abdominal cavity of mice was promptly opened, and the hepatic portal vein was located after anesthesia. A soft needle was inserted for perfusion with Perfusion buffer (100mL 1× HBSS, 200uL 0.5mol/L EGTA) for 3-5 minutes, while the inferior vena cava was cut to clear the liver. Subsequently, Digestion buffer (60mL 1× HBSS, 120uL 2M $MgSO_4$, 60uL 1.25M $Cacl_2$•$H_2O$, 0.015g Collagenase I) was injected at a consistent rate for approximately 15 minutes. Following perfusion, the liver was delicately transferred into a pre-cooled petri dish with PBS, cut into 2-3 mm fragments, and incubated at 37℃ for 15 minutes in Digestion buffer. The liver tissue fluid was then filtered through a 70um cell filter and centrifuged at 50g for 5 minutes (ac/brake=0) at 4℃ to collect the supernatant. This supernatant underwent centrifugation at 450g for 5 minutes (ac/brake=5) at 4℃ to obtain the NPCs population. The NPCs were suspended in a 15mL centrifuge tube with 4mL 20% Optiprep solution, followed by centrifugation at 3000 rpm for 17 minutes (ac/brake=0) at 4℃. After centrifugation, the cells at the boundary between the Optiprep solution and 1xHBSS buffer were collected into a new tube, supplemented with 1xHBSS buffer, and centrifuged for 5 minutes at 450g (ac/brake=5). Red blood cells (RBCs) were lysed with 1mL ACK buffer (150mM NH4Cl, 10mM KHCO3, 0.1mM Na2EDTA, pH=7.2-7.4), neutralized with 1xHBSS buffer, and centrifuged. The cells were resuspended in cell medium and placed in a 3.5cm petri dish for 10 minutes to obtain KCs by washing out un-adherent cells with PBS. The purity and viability of isolated KCs reached 90% and 80%, respectively, which was confirmed by staining with anti-Timd4 antibodies or trypan blue.

**Preparation of Bone Marrow Derived Macrophages**

To obtain bone marrow-derived macrophages, femur and tibia bone marrow from healthy male *C57BL6/J* was extracted, resuspended in DMEM/F12 medium (VivaCell, Cat# C3113-0500) containing 10% FBS (VivaCell, Cat# C04001-500) and 20% L929 conditioned medium, seeded in culture dishes and cultured at 37 ℃ with 5% $CO_2$ in a humidified atmosphere for 7 days. Fresh medium was added on day 4. The cells were maintained in a standard 37 ℃ in 5% $CO_2$ incubator.

**Flow Cytometry**

The NPCs were resuspended in fluorescence-activated cell sorting (FACS) buffer, consisting of PBS supplemented with 2% bovine serum albumin. 1uL of anti-CD16/CD32 antibody (Invitrogen, Cat# 14-0161-86) was added to the cell suspension, and the mixture was incubated at 4℃ for 10 min to block any nonspecific binding. After blocking, the Mouse NPCs were labeled with monoclonal antibodies conjugated with fluorescent dyes. This labeling process was carried out at 4℃ for 30 min. The labeled cells were washed thrice with cold FACS buffer. The antibodies used for labeling were as follows: anti-mouse F4/80(APC) (Invitrogen, Cat# 17-4801-82), anti-CD45 (eFluor450) (Invitrogen, Cat# 48-0451-82), anti-mouse Tim-4(PE) (Invitrogen, Cat# 12-5866-82), and anti-mouse CD11b (PerCP/Cyanine5.5) (BioLegend, Cat# 101228). All the primary antibodies were used at a dilution of 1:100. The samples were analyzed by flow cytometry using an LSR Fortessa Cell Analyzer (BD Biosciences). The data obtained were further analyzed using FlowJo version 10.0.

**Histology & immunofluorescence**

*H&E staining* Tissues were fixed with buffered 10% paraformaldehyde (Sangon Biotech, Cat# A500684-0500) overnight at 4°C and embedded in paraffin. Ultra-thin tissue slices (5μm) were prepared and deparaffinized. H&E staining was performed on the tissue sections, and the slides were examined under a microscope (Olympus, BP80).

*Immunofluorescence on frozen section* Immunofluorescence staining was conducted on frozen sections of fresh liver tissues. Initially, the tissues were fixed using 2% paraformaldehyde for 1 h and subsequently dehydrated overnight in a 30% sucrose solution. The following day, tissue embedding was performed using OCT (SAKURA, Cat# 4583), after which ultra-thin sections of 5μm were sliced. Permeabilization was achieved using 0.02% Triton X-100 for 10 minutes at room temperature, followed by blocking with 5% normal goat serum (VivaCell，Cat# C2530-0100). Primary antibodies against mouse F4/80 (Biolegend， Cat# 123140, 1:300, Alexa Fluor 594), Timd4 (Biolegend， Cat# 130008, 1:300, Alexa Fluor 647), and Chi3l1 (Abcam, ab180569, 1:400) were then applied. Secondary antibodies (Affinipure Goat Anti-Rabbit 488-conjugated, Jackson ImmunoResearch, 111-545-003, 1:1000 and Goat Anti-Mouse 594-conjugated, Jackson ImmunoResearch, 115-585-003, 1:1000) were used accordingly, with F4/80 and Timd4 antibodies being self-fluorescent and thus not requiring secondary labeling. After washing, the slides were mounted using antifade medium, and nuclei were stained with DAPI. Additionally, Tunel staining was performed on separate frozen sections of liver tissues. Following fixation with 4% paraformaldehyde and treatment with Proteinase K (20 μg/mL in PBS, BBI, Cat# B600169-0002), the sections were permeabilized with Triton X-100 and blocked with normal goat serum. Subsequently, the sections were incubated with an anti-mouse Clec4f antibody (BioLegend， Cat# 156804, 1:200), followed by Tunel staining (servicebio, Cat# G1502-100T) according to the manufacturer's instructions. Nuclei were counterstained with DAPI, and images were captured using a confocal laser scanning microscope (ZEISS, LSM900).

*Oil Red O staining* Oil red O staining was conducted on unfixed frozen sections embedded directly in OCT. Frozen sections of the liver were cut at a thickness of 10 μm. After rinsing with water, the sections were immersed in 60% isopropanol for 2 min. Subsequently, the sections were stained with an Oil Red O staining solution (Solarbio, Cat# IO1720) at 37°C for 10 to 15 min. Following staining, the sections were immediately placed in 60% isopropanol and washed to 3-5 times to eliminate excess dye solution. Nuclei were counterstained with a hematoxylin staining solution. After rinsing with distilled water, the sections were sealed with glycerol gelatin (Solarbio, Cat# S2150).

*Sirius red staining* Tissues were fixed with buffered 10% paraformaldehyde (Sangon Biotech, Cat# A500684-0500) overnight at 4°C and embedded in paraffin. Ultra-thin tissue slices (5μm) were prepared and deparaffinized. Sirius red staining (Solarbio, Cat#G1472) was performed according to the manufacturer's instructions, and the slides were examined under a microscope (Olympus, BP80).

*Immunofluorescent staining on BMDM or KCs* Cells were seeded on coverslips in 12-well plates and treated with or without recombinant murine Chi3l1(rChi3l1, 100ng/ml, SB, Cat# 50929-M08H; diluted in PBS) under no glucose or high glucose (25mM, VivaCell, Cat# C3113-0500) for 24 h. The cell slides were washed with cold PBS and fixed with 4% paraformaldehyde in PBS for 10 min at room temperature. The cells were permeabilized with 0.02% Triton X-100 for 10 min at room temperature. After blocking with 5% normal goat serum, cells were incubated with primarily antibodies anti-STBD1 (Proteintech, Cat# 11842-1-AP, 1:300) overnight at 4℃ to label glycogen in the cells. On the second day, after washing with 0.05% PBST, cells were incubating with 594-conjugated Goat Anti-Mouse IgG (H+L) (Jackson ImmunoResearch, 115-585-003, 1:1000) for 1 h at room temperature. After rinsing with 0.05% PBST, cells were counterstained with DAPI (beyotime, C1006) and mounted onto slides. Images were captured with an Olympus BP80 microscope. The immunofluorescent staining on KCs follows a similar protocol to that used for BMDM. Primary antibodies against Timd4 (Biolegend, Cat# 130008, 1:300, Alexa Fluor 647) are employed to label KCs.

*Utilize 2-NBDG to track glucose uptake* To assess glucose uptake, 2-NBDG, a fluorescent glucose derivative, was utilized as a substitute for glucose in detecting glycogen formation within live cells.[3] Following 12 h of glucose deprivation, cells were treated with 20 μM 2-NBDG (Invitrogen, Cat# N13195; diluted in PBS) for 6 h. Post-treatment, cells were fixed with paraformaldehyde and counterstained with DAPI before being mounted onto slides using coverslips. Imaging of the samples was performed using a confocal laser scanning microscope (ZEISS, LSM900), facilitating visualization and analysis of glucose uptake dynamics.

*Calcein/PI staining* Live cell staining using Calcein/PI was conducted following the manufacturer's instructions from a commercially available kit (Beyotime, Cat# C2015M). Isolated KCs were seeded in 12-well plates and subjected to various treatments. In one set of experiments, cells were treated with either Isopropyl alcohol (1μl, Sangon Biotech, Cat# A507048-0500) as a control or palmitic acid (800mM in 1μl, Sigma, Cat# P0500), or 100ng/ml rChi3l1 with palmitic acid, all for 24 h.

**Western blot analysis**

The sample preparation and procedure were conducted as previously described.[1] The following antibodies were used: anti-Caspase 3 (Cell Signaling Technology, Cat# 9662S, 1:1000), anti-Cleaved Caspase-3 (Cell Signaling Technology, Cat# 9664S, 1:1000), anti-β-actin (Proteintech， Cat# 66009-1-lg, 1:1000), anti-Chi3l1 (Proteintech, Cat# 21829-1-AP, 1:1000), anti-Albumin (Proteintech, Cat# 21829-1-AP, 1:1000), anti-GAPDH (CWBIO, Cat# cw0100M, 1:1500), anti-α-SMA (Invitrogen, Cat# 50-9760-82, 1:1000). Peroxidase-conjugated Affinipure Goat Anti-Mouse IgG(H+L) (Jackson ImmunoResearch, 115-035-003, 1:2000), Peroxidase-conjugated Affinipure Goat Anti-Rabbit IgG(H+L) (Jackson ImmunoResearch, 111-035-003, 1:2000) and Peroxidase-conjugated Affinipure Goat Anti-Rat IgG(H+L) (Jackson ImmunoResearch, 112-035-003, 1:2000) were used for secondary antibody incubation.

**Microscale Thermophoresis (MST) Assay**

The dissociation constant (Kd) for the interaction between Chi3l1 and glucose was determined using a Microscale Thermophoresis (MST) instrument (NanoTemper Technologies). Mouse Chi3l1 protein was labeled with a His-Tag Labeling Kit RED-tris-NTA 2nd Generation (NanoTemper Technologies, Cat# MO-L018) following the manufacturer's instructions. Briefly, 90 μL of dye solution (final concentration 100 nM) was mixed with 90 μL of purified Chi3l1 protein and incubated at room temperature for 30 min. The mixture was then centrifuged at 15,000 × g for 10 min at 4 °C, and the supernatant was collected for subsequent binding assays. Serial dilutions of glucose were prepared in 1× PBS-T buffer (1× PBS containing 0.05% Tween-20, pH 7.4) to generate 16 concentration gradients. Equal volumes of labeled Chi3l1 protein were added to the diluted glucose solutions and incubated at room temperature in the dark for 30 min. The samples were loaded into premium capillaries (NanoTemper Technologies, Cat# MO-K022), and MST measurements were performed at 25 °C using medium MST power and 100% excitation power. Data were collected and analyzed using MO.Affinity Analysis Software (version 2.3) to calculate the dissociation constant (Kd).

**Biotin-glucose pull-down assay**

Biotin-glucose chemical synthesis and biotin-glucose pull-down assays were conducted according to the established protocols.[4] In summary, Magic Dynabeads M-280 Streptavidin (Invitrogen, Cat# 11205D) was pre-incubated with either free biotin or biotin-labeled glucose at 4℃ for 1 h. Next, the beads were incubated with serum from mice fed HFHC diet for 16 weeks overnight at 4℃. Afterward, the beads were washed six times with wash buffer and subjected to immunoblotting.

**Preparation of single-cell RNA-seq library and data analysis**

After isolating single NPCs following the procedure described above, the cells were captured using the BD Rhapsody system (BD Biosciences, Cat# 633731). Subsequently, the captured single cells underwent whole transcriptome amplification following the BD Rhapsody workflow (BD Biosciences, Cat# 633733). Transcriptome libraries were then prepared from the amplified cDNA using the library preparation protocols provided by BD Rhapsody (BD Biosciences, Cat# 633801). Quality control checks (Revvity) were conducted on the prepared libraries to ensure proper amplification and library construction. Finally, the libraries were sequenced using BD platforms.

The scRNA-seq data were processed and analyzed using R (version 4.0.5) and the R/Seurat package (version 4.2.3). To ensure data quality, low-quality cells, empty droplets, and cells from multiplexed captures were excluded based on the distribution of unique genes detected in each cell. Additionally, genes expressed in fewer than three cells in a sample were excluded. Cells with fewer than 200 genes or more than 4000 genes were excluded, and cells with a mitochondrial gene fraction exceeding 25% were removed. After quality control, datasets from different batches were normalized using SCTransform. For the integrated data, reciprocal PCA-based integration was performed using the FindIntegrationAnchors and IntegrateData functions. Subsequent clustering was conducted using the FindClusters function with resolution values of 0.8 or 1.0, depending on the dataset. Normalization accounted for total unique molecular identifier (UMI) and mitochondrial gene content, and analysis was performed using the 4000 most variable genes. Visualization of the clustered data was achieved using Uniform Manifold Approximation and Projection (UMAP).

To identify cluster biomarkers, we utilized the Find Markers function and Find All Markers function in Seurat, setting a minimum percentage of expression (min.pct) of 0.1 and a log-fold change threshold (logfc.threshold) of 0.25. Cell types were annotated using marker genes with CellMaker 2.0. Subsequently, remaining cells associated with KCs and monocytes were extracted and reclustered. We conducted re-normalization, RPCA-based integration, and cluster stability analysis, determining an optimal resolution of 0.5 or 0.3, depending on the dataset. Cell annotations were based on known cell-lineage-specific marker genes. [5] For differential gene and pathway analysis, differential expression analysis between groups such as NCD and HFHC or NCD, WT HFHC, and *Chil1*$^{-/-}$ HFHC was performed using the Find Markers function. Pathway analysis utilized the Kyoto Encyclopedia of Genes and Genomes (KEGG), while gene ontology (GO) analysis was conducted using the cluster Profiler package to explore functional categories associated with differentially expressed genes. The most highly enriched pathways were identified using the enrich GO and enrich KEGG functions from the cluster Profiler package. Additionally, Gene Set Variation Analysis (GSVA) was employed to explore associations between cell death and other pathways, utilizing the gsva function from the GSVA R package. Pathways were visualized using the pheatmap R package.

**Extracellular acidification rate (ECAR) measurement**

The ECAR measurements were conducted according to the manufacturer's instructions (Agilent, Cat# 103020-100). Briefly, mature BMDM were cultured in a Seahorse XF24 cell culture plate. After 12 h of culture, the DMEM culture medium was pre-treated for 24 h. 1 h before the analysis, the culture medium was changed to the corresponding XF basal medium (Agilent, Cat#103334-100) supplemented with glutamine (The final concentration was 2mmol/L, Agilent, Cat#103579-100), and the culture plates were incubated at 37℃ without $CO_2$. Compounds were added in the following order: 10mM glucose, 1.0 mM oligomycin, and 50mM 2-deoxyglucose. Measurements were conducted using a Seahorse XF24 analyzer (Agilent Technologies). After completing the SeahorseXF glycolytic stress test, basic glycolytic ability and total glycolytic ability were calculated based on the generated report.

**RNA Extraction and Quantitative Real-Time PCR**

Total RNA was extracted from cultured cells or ground tissues using TRIzol reagent (Invitrogen, Cat# 15596018) according to the manufacturer's instructions. Briefly, lysates were centrifuged at 12,000 × g for 10 min at 4 °C, and the supernatant was transferred to a new tube. Chloroform (200 µL) was added, mixed thoroughly, and incubated at room temperature for 10 min, followed by centrifugation. The aqueous phase was collected, mixed with an equal volume of isopropanol, and incubated for 10 min at 4 °C to precipitate RNA. After centrifugation, the pellet was washed twice with 75% ethanol, air-dried for 10 min at room temperature, and dissolved in RNase-free water. RNA concentration and purity were assessed using a NanoDrop spectrophotometer (Thermo Fisher Scientific). Complementary DNA (cDNA) was synthesized from total RNA using the PrimeScript RT Reagent Kit (Takara, Cat# 6210B) following the manufacturer's instructions. Quantitative real-time PCR (qPCR) was performed using SYBR Green Master Mix (Thermo Fisher, Cat# A25742) in triplicate on a QuantStudio 1 Real-Time PCR System (Life Technologies, Grand Island, NY, USA). Relative gene expression was calculated using the ΔΔCt method, and primer sequences are listed in Table S2.

**Table S2**
**qPCR primers**

| Targeted gene | Primer | Sequence |
|---|---|---|
| musChil1 | F(5'-3') | AGAAACACCAACCTGAAGACC |
| | R(5'-3') | CCCATCAAAGCCATAAGAACG |

**In Vivo Glucose Uptake Assay**

In vivo glucose uptake was assessed as previously described[6]. Mice were fasted for 16 h and subsequently refed. At 45 min after refeeding, mice were intraperitoneally injected with 2-NBDG (100 µL, 12 mg/kg body weight). Mice were euthanized 10 min post-injection, and liver tissues were collected for analysis. For quantification of total glucose uptake, weighed portions of liver tissue were homogenized in cell lysis buffer (150 mM NaCl, 50 mM HEPES, pH 7.4, 1% Triton X-100). After centrifugation, 2-NBDG fluorescence intensity in the supernatant was measured using a microplate reader (excitation/emission: 465/540 nm). For histological analysis, liver samples were dehydrated overnight in 30% sucrose, embedded in OCT compound (SAKURA, Cat# 4583), and sectioned at 5 µm thickness. Sections were permeabilized with 0.02% Triton X-100 for 10 min at room temperature and blocked with 5% normal goat serum (VivaCell, Cat# C2530-0100). Kupffer cells were identified using an anti-Timd4 antibody (BioLegend, Cat# 130008; 1:300, Alexa Fluor 647). Nuclei were counterstained with DAPI, and images were acquired using a confocal laser scanning microscope (ZEISS LSM900).

**Data analysis from Gene Expression Omnibus (GEO) database**

The mRNA expression of Chi3l1 was analyzed using liver tissue sequencing data obtained from the NCBI Gene Expression Omnibus database (www.ncbi.nlm.nih.gov/geo). The dataset consisted of global RNA sequencing results from 51 patients with MASLD and 47 patients with MASH. The accession number for this dataset was GSE167523. Additionally, RNA sequencing results from the same database were obtained for liver tissues from 5 patients without MASLD, 15 patients with MASLD, and 10 patients with MASH (accession number: GSE207310). Liver tissue sequencing data from 78 patients with MASLD were obtained from a database. This included 20 patients without MASH and 58 patients with MASH (accession number: GSE130970).

**Statistical analysis**

Data are presented as mean ± standard error of the mean (SEM) in all graph figures. Statistical analyses were conducted using the SPSS statistics software (Version 22). To compare the two groups, an unpaired two-tailed Student's t-test was used. One-way analysis of variance (ANOVA) was performed for comparisons involving three or more groups. For patients with non-alcoholic fatty liver, the samples were tested using the Mann-Whitney test. Statistical significance was set at $p < 0.05$ and p value is indicated. All cell culture results represent at least three independent experiments.

**Reference**


1 Yan Chen, Bin Qi, Zhao Shan. Peptidoglycan-Chi3l1 interaction shapes gut microbiota in intestinal mucus layer. *Elife* **13**, doi:https://doi.org/10.7554/eLife.92994.1 (2024).

2 Shan Z, Li L, Atkins CL, *et al.* Chitinase 3-like-1 contributes to acetaminophen-induced liver injury by promoting hepatic platelet recruitment. *eLife* **10**, doi:10.7554/eLife.68571 (2021).

3 Liu QX, Li JX, Zhang WJ, *et al.* Glycogen accumulation and phase separation drives liver tumor initiation. *Cell* **184**, 5559, doi:10.1016/j.cell.2021.10.001 (2021).

4 Chen TJ, Xu ZG, Luo J, *et al.* NSUN2 is a glucose sensor suppressing cGAS/STING to maintain tumorigenesis and immunotherapy resistance. *Cell Metabolism* **35**, 1782-+, doi:10.1016/j.cmet.2023.07.009 (2023).

5 Li L, Cui L, Lin P, *et al.* Kupffer-cell-derived IL-6 is repurposed for hepatocyte dedifferentiation via activating progenitor genes from injury-specific enhancers. *Cell Stem Cell* **30**, 283, doi:10.1016/j.stem.2023.01.009 (2023).

6 Shi X, Hu X, Fang X, *et al.* A feeding-induced myokine modulates glucose homeostasis. *Nat Metab* **7**, 68-83, doi:10.1038/s42255-024-01175-9 (2025).
